# Putative Looping Factor ZNF143/ZFP143 is an Essential Transcriptional Regulator with No Looping Function

**DOI:** 10.1101/2024.03.08.583987

**Authors:** Domenic N. Narducci, Anders S. Hansen

## Abstract

Interactions between distal loci, including those involving enhancers and promoters, are a central mechanism of gene regulation in mammals, yet the protein regulators of these interactions remain largely undetermined. The zinc finger transcription factor ZNF143/ZFP143 has been strongly implicated as a regulator of chromatin interactions, functioning either with or without CTCF. However, ZNF143/ZFP143’s role in this process and its function, either with or without CTCF, are not well understood. Here, we tagged both CTCF and ZNF143/ZFP143 with dual-purpose degron/imaging tags to combinatorially assess their looping function and effect on each other. We find that ZNF143/ZFP143 possesses no general looping function in mouse and human cells, and that it largely functions independently of CTCF. Instead, ZNF143/ZFP143 is an essential and highly conserved transcription factor possessing an extremely stable chromatin residence time (>20 min) that regulates an important subset of mitochondrial and ribosomal genes.

## INTRODUCTION

Precise control of gene expression is important for nearly all processes within cells and tissues. In mammals, gene expression control is tightly regulated by looping interactions between enhancers and promoters^1–3^, and dysregulation of this process can lead to developmental defects and disease^4–10^. Therefore, understanding the way proteins in the nucleus regulate these long-range looping interactions is a vital biological question. These interactions are constrained by topologically associating domains (TADs), genomic regions enriched for self-interaction. TADs are formed when the loop extrusion factor cohesin stalls at CTCF boundaries, promoting more frequent enhancer-promoter (E-P) and promoter-promoter (P-P) interactions within TADs than between across TAD boundaries^1,8,11–19^. However, because CTCF is expressed in all cell types, it is currently unclear how cell-type specific 3D genome structure is regulated^14,20,21^. Furthermore, acute depletion of CTCF causes dysregulation of only a few hundred genes and a small fraction of E-P/P-P interactions^21–24^. Together, these results suggest the existence of other proteins that regulate loop and domain formation.

Here, we focus on the seven C2H2 zinc finger protein ZFP143/ZNF143, which has been implicated as a regulator of loop and domain formation due to its very strong enrichment at loop anchors, including both structural and E-P loops^17,25–40^. Though ZFP143 refers to the mouse protein and ZNF143 to the human protein, in this study, we will refer to both as ZFP143 for simplicity. ZFP143 was initially discovered as the *Xenopus laevis* gene *staf*, a transcriptional regulator^41–45^ that binds to promoters in a sequence-specific manner and drives gene expression^46–53^. It was found that aberrant expression of ZFP143 modulates proliferation, migration, and the cell cycle in cancer^54–64^ and contributes to chemotherapeutic resistance through modulation of DNA damage repair^65,66^, and that mutations in ZFP143 or in its binding sites can cause metabolic disease and endothelial corneal dystrophy^26,67,68^.

ZFP143 was first suggested to regulate chromatin looping based off analysis of ChIP-seq data from the ENCODE consortium^33,34^. It was reported that ZFP143 cobinds alongside known looping regulators CTCF and cohesin^33–35,69–71^, and Hi-C and Micro-C studies found ZFP143 to be one of the most highly enriched proteins at domain boundaries^17,32^, a result which held in several computational analyses aimed at architectural proteins^36– 38,70^. It was additionally reported that ZFP143 participates in chromatin interactions involving promoters^25,28^, and that siRNA-mediated knockdown of ZFP143 showed a general reduction in loop strength measured by Hi-C^30^. Deletion or siRNA-mediated knockdown of ZFP143 appears to show specific weakening of short-scale E-P loops^27–29^, and several studies have proposed mechanisms involving direct interaction with or recruitment by CTCF to explain these changes in looping^27,29^. However, because no study has acutely perturbed ZFP143 in these contexts, it remains to be elucidated whether ZFP143 directly regulates looping or CTCF binding.

Here, we acutely deplete both ZFP143 and CTCF on short time scales and evaluate their role in directly regulating looping using Micro-C, a method uniquely suited to resolving fine-scale interactions^23,32,72–74^. We find that loss of ZFP143 has no effect on looping. Additionally, we evaluate the impact of ZFP143 binding on CTCF and *vice versa* using a combination of ChIP-seq and live-cell imaging. We find that ZFP143 binds largely independently of CTCF, but, like CTCF, it has a very long chromatin residence time compared to other transcription factors^75^. Lastly, we use acute perturbation of ZFP143 to examine its role as a transcriptional regulator and find that ZFP143 regulates an important subset of housekeeping genes. Taken together, these results indicate that ZFP143 does not function as a regulator of chromatin architecture, with or without CTCF, but rather, it stably binds chromatin and acts as an essential transcriptional regulator.

## RESULTS

### ZFP143 is an essential protein that stably binds chromatin

To investigate ZFP143’s role in chromatin looping and its relationship to CTCF in mouse embryonic stem cells (mESCs), we tagged both ZFP143 and CTCF with orthogonal, multi-purpose tags consisting of an mAID2^76^ or FKBP12(F36V)^77^ degron, a Halo-or SNAP_f_-tag, and an epitope tag using CRISPR/Cas9-mediated genome editing (**Fig. 1A**). To control for clone-to-clone variation, we independently established two tagged cell lines (clones A and B) as well as a third mESC line with the FKBP12(F36V) and SNAP_f_-tag on ZFP143 instead of CTCF (clone D). We also acquired a previously established human HEK293T ZNF143 degron cell line (clone 30)^53^ to assess inter-species effects. We validated homozygous tagging by PCR and near wild-type expression levels of both ZFP143 and CTCF by western blot and confirmed nuclear localization by live-cell imaging (**Fig. 1B-C**). Imaging of ZFP143 revealed a somewhat punctate appearance similar to CTCF^12,78–80^ (**Fig. 1C**). To validate the degron tags, we performed depletion timecourses and determined three hours to be an optimal timepoint since it minimizes secondary effects and is the earliest time where both ZFP143 and CTCF were completely degraded (**Fig. 1B, Fig. S1A**). We also confirmed this timepoint in clones D and 30 (**Fig. 1B**). Finally, we asked whether ZFP143 is essential in mESCs. We grew clone D mESCs under ΔZFP143 conditions for six days and found their growth rate was strongly impaired (**Fig. 1D, Fig. S1B**). Next, we attempted to generate a ZFP143 knockout cell line using CRISPR/Cas9-mediated genome editing which produced 39% heterozygous and 0% homozygous deletions in 191 screened colonies suggesting mESCs are not viable with a homozygous ZFP143 deletion (**Fig. S1C**). Taken together, these results show ZFP143 is essential in mESCs consistent with prior work^27,48,53,81,82^.

**Figure 1.**
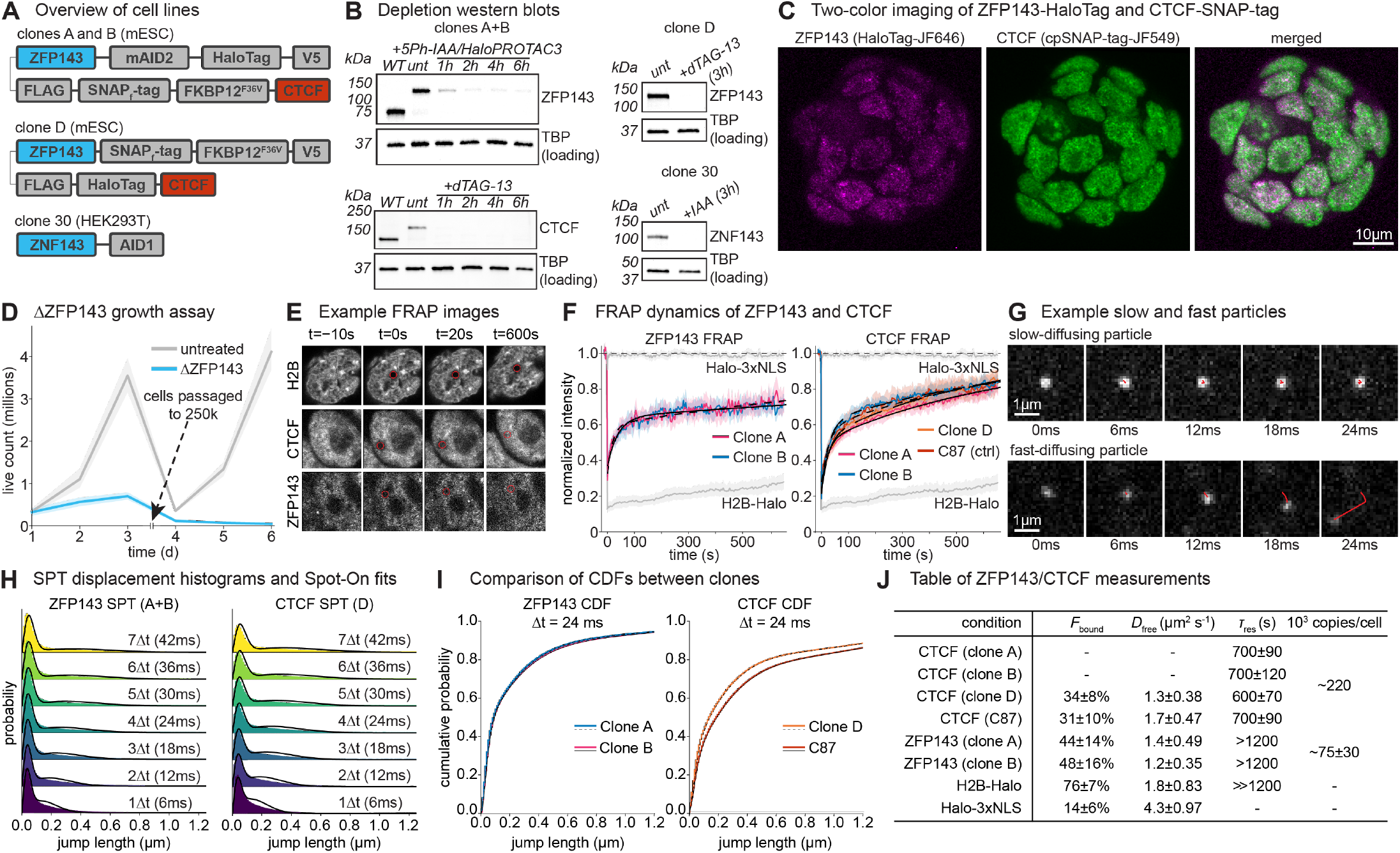
ZFP143 can be homozygously tagged and stably binds to chromatin. (**A)** Overview of the genome engineered cell lines. (**B**) Western blots showing depletions of either ZFP143 or CTCF for all clones. For clones A and B (left), ZFP143 (top) shows the expected size shift of 48.8 kDa and near complete degradation after two hours. CTCF (bottom) likewise shows the expected size shift of 37.2 kDa and near complete degradation after one hour. Clones D and 30^53^ (right) show near complete degradation at the selected three-hour timepoint. TBP was used as a loading control in all western blots. (**C**) Live-cell imaging of a single clone A mouse colony validating the fluorescent labeling approach and showing nuclear localization of both CTCF and ZFP143. (**D**) Growth curve showing live cell counts in clone D with (blue) and without (gray) constitutive ZFP143 depletion for six days. After counting on day three, cells were reseeded at a density of 250k live cells per well. Error bars indicate 95% confidence intervals. (**E**) Example images of FRAP recovery in the H2B-Halo, Halo-CTCF, and ZFP143-Halo conditions at 10 seconds prior to the bleach, on the frame of the bleach, 20 seconds after bleaching, and 600 seconds after bleaching. (**F**) FRAP recovery curves showing normalized intensity of ZFP143 recovery (left) for clones A (magenta) and B (blue) over time and CTCF recovery (right) for clones A (magenta), B (blue), and D (orange) as well as the C87 Halo-CTCF line^12^ (red). Halo-NLS and H2B-Halo control curves are shown in gray. Error bars indicate 95% confidence intervals. (**G**) Example images of particles observed in single particle tracking experiments. (**H**) Jump displacement histograms for CTCF (clone D; left) and ZFP143 (clones A and B; right) showing jump length probabilities for seven different time lags. (**I**) Cumulative distribution functions (CDFs) comparing jump length probabilities between clones A and B (left) and clone D and C87 (right). (**J**) Table summarizing the key measurements plus or minus one standard deviation from SPT, FRAP, and absolute abundance quantification.

The established loop regulator CTCF exhibits an unusually long residence time and high bound fraction^12,75^. To understand if ZFP143 exhibits similarly stable binding, we performed live-imaging. First, we determined ZFP143’s residence time using fluorescence recovery after photobleaching (FRAP). By labeling cells with Halo-JFX554 or cpSNAP-JFX554^83^ and measuring the fluorescent intensity inside a 1 µm circle following photobleaching, we were able to fit a previously derived reaction-dominant model^84^ (**Fig. 1E-F**). Since the fitted residence time was much longer than the time of the experiment (∼10 min), we were unable to precisely determine it, but we estimate a minimal residence time of at least ∼20 minutes. This was longer even than the ∼600-700s residence time we measured for CTCF. Next, to determine the chromatin-bound fraction of ZFP143 and CTCF, we performed single-particle tracking (SPT). Here, we used a combination of highly inclined laminated optical sheet illumination^85^, high laser power, and ∼6 ms exposure times with 1 ms stroboscopic excitation to follow single molecules of ZFP143 or CTCF as they diffused within the nucleus^86^ (**Fig. 1G**). We calculated jump displacement histograms using subpixel localizations of tracked particles and fit the previously established two-state Spot-On model^86^ (**Fig. 1H-I, Fig. S1D-E**). We found that ZFP143 has a bound fraction between 40 and 50% and a free diffusion coefficient on the order of 1.3 µm^2^ s^-1^. Furthermore, we found ZFP143’s chromatin binding and long residence time to require its seven zinc-finger DNA-binding domain (**Fig. S1F**). Lastly, we measured that mESCs have on average ∼ 75,000 ZFP143 copies per cell (**Fig. 1J, Fig. S1G**). Together, these results suggest a model where ZFP143 interacts with its cognate binding sites in a DNA binding domain (DBD)-dependent manner with, to our knowledge, the longest residence time currently measured for a mammalian transcription factor.

### ZFP143 is not a regulator of chromatin looping

To evaluate ZFP143’s putative looping function, we performed Micro-C^73,87^ following three-hour depletions of either ZFP143, CTCF, or both ZFP143 and CTCF and found high reproducibility between all replicates and clones A and B, achieving between ∼700 million and 1 billion unique contacts per condition (**Fig. S2A**). To evaluate global effects on interaction probability we generated *P(s)* curves for all conditions and found no change upon ZFP143 depletion (**Fig. S2B**). Likewise, we observed no substantial effects on compartments (**Fig. S2C-D**) nor TADs in terms of insulation score and strength (**Fig. S2E-G**). To identify loops anchored by ZFP143 and CTCF, we performed ZFP143 and CTCF ChIP-seq in untreated cells as well as all three depletion conditions. Likewise, we performed precision run on sequencing (PRO-seq)^88,89^, a nascent transcriptomics method capable of resolving changes on the timescale of the depletions, in all conditions to evaluate potential impacts of these interactions on gene expression. Even at sites where loops are anchored by ZFP143 binding on at least one side, Micro-C maps are largely invariant to ZFP143 depletion (**Fig. 2A-B**). In contrast, loops and domains are largely eliminated by CTCF depletion as previously described^22,23^. Conversely, gene expression at ZFP143-bound sites is often dependent on ZFP143 binding and independent of local, CTCF-driven structure. For example, at both the *Timm13/Lmnb2* locus and the *Rpp30 locus*, “dots” corresponding to looping interactions are preserved upon ZFP143 depletion, even at loops lining up with a ZFP143 ChIP-seq peak, while both the CTCF depletion and double depletion contact matrices are largely devoid of “dots”/loops (**Fig. 2A-B**). Although ZFP143 anchored loops were unaffected, ZFP143-bound genes changed expression as measured by PRO-seq. *Timm13* and *Lmnb2* PRO-seq signal decreased ∼3.2-fold and ∼4.9-fold, respectively (**Fig. 2A**), while *Rpp30* signal is likewise reduced by a factor of ∼4.9 (**Fig. 2B**). Aggregate peak analysis (APA) using loops from Hsieh *et al*. 2020 revealed neither dependence on ZFP143 nor redundancy between ZFP143 and CTCF (**Fig. 2C**). Likewise, ZFP143 depletion had no effect on cohesin-bound, P-P, E-P, and ZFP143-bound loops. In fact, CTCF depletion had a greater effect than ZFP143 depletion on all loop subsets including even ZFP143-bound loops. To ensure this observation was not due to the existence of two oppositely affected loop populations, we plotted loop strengths on a per-loop basis in all depletion backgrounds and found no such effects (**Fig. 2D**). Due to the surprising inconsistency with ZFP143’s reported looping function^17,25–32,34–38^, we repeated our experiments in clone D which had a more complete ZFP143 depletion than clones A and B (**Fig. 1B**). We again found that even at sites where ZFP143 is bound adjacent to a loop, there is no effect on global (**Fig. S2B-D**) or local 3D structure (**Fig. 2E-F, Fig. S2F**,**H**,**I**). Moreover, APA again revealed no dependence on ZFP143 in any of the loop classes (**Fig. 2G**). Lastly, to test for effects on insulation that were independent of loop strengths, we performed metaplots over boundaries in insulation score tracks and again found no ZFP143-mediated effects nor interactions between ZFP143 and CTCF (**Fig. 2I**). We conclude that ZFP143 is not a regulator of chromatin looping in mESCs.

**Figure 2.**
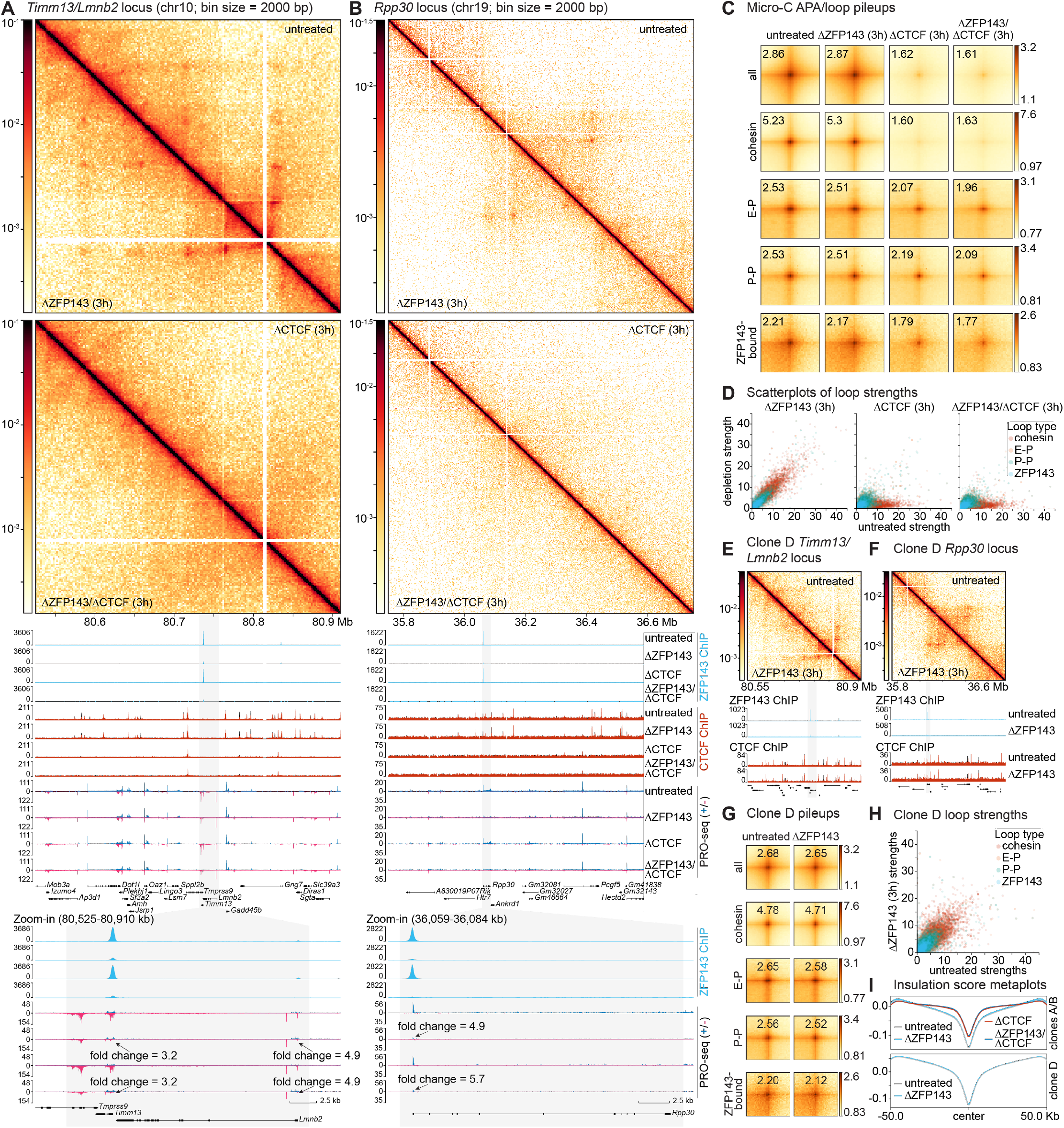
Loops are unaffected by ZFP143 depletion in mESCs. (**A-B**) Panels showing genomics data at two representative loci averaging clones A and B. Shown from top to bottom are Micro-C maps at 2000 bp resolution comparing untreated to ΔZFP143 (3h), maps comparing ΔCTCF (3h) to ΔZFP143/ΔCTCF (3h) double depletion, ZFP143 ChIP-seq tracks (cyan), CTCF ChIP-seq tracks (red), and PRO-seq with plus strand in blue and minus strand in magenta for untreated, ΔZFP143 (3h), ΔCTCF (3h), and ΔZFP143/ΔCTCF (3h) double depletion conditions. Shown below, are zoomins on ZFP143-regulated genes showing reduction in PRO-seq signal upon ZFP143 depletion. (**A**) Shown is the *Timm13*/*Lmnb2* locus. (**B**) Shown is the *Rpp30* locus. (**C**) Aggregate peak analysis (APA)/loop pileup analysis for all called loops, cohesin-bound loops, E-P loops, and P-P loops from Hsieh *et al*. 2020 and ZFP143-bound loops across untreated, ΔZFP143 (3h), ΔCTCF (3h), and ΔZFP143/ΔCTCF (3h) depletion conditions in log_10_ scale. Shown in the upper left corner is the average signal within the plot. (**D**) Scatterplots of loop strengths calculated as in (**c**) for cohesin loops (red), E-P loops (orange), P-P loops (teal), and ZFP143 loops (cyan) comparing either ΔZFP143 (3h), ΔCTCF (3h), or ΔZFP143/ΔCTCF (3h) to untreated. (**E-F**) Panels showing genomics data at the two representative loci in (**A-B**) in clone D. Shown from top to bottom are Micro-C maps at 2000bp resolution comparing untreated to ΔZFP143 (3h), ZFP143 ChIP-seq tracks (cyan), and CTCF ChIP-seq tracks (red) for untreated and ΔZFP143 (3h) depletion conditions. (**G**) APA/loop pileup analysis as in (**c**) repeated for clone D. (**H**) Scatterplots of boundary strengths as in (**D**) repeated for clone D comparing ΔZFP143 (3h) to untreated conditions. (**I**) Metaplots over insulation score tracks called at 50 kb resolution comparing untreated (gray), ΔZFP143 (3h) (cyan), ΔCTCF (3h) (red), and ΔZFP143/ΔCTCF (blue) double depletion (3h) in clones A and B (top) and clone D (bottom).

Although our results rule out a looping function for ZFP143 in mESCs, prior work proposing a looping function for ZFP143 have come from both mouse and human cell lines. Therefore, to test if ZFP143 regulates chromatin loops in human cells, we acquired a previously validated HEK293T ZFP143-degron cell line^53^ (**Fig. 1A**, clone 30) and performed Micro-C following a three-hour depletion of ZFP143 with strong reproducibility between the two replicates, achieving ∼1 billion unique contacts per condition (**Fig. S3A**). We again found no impact on *P(s)* curves, compartments, and insulation scores (**Fig. S3B-G**). Like in mESCs (**Fig. 2**), at the *ABL1* locus, a strong ZFP143-anchored loop was unaffected by ZFP143 depletion (**Fig. 3A**). Despite not affecting loops, ZFP143 depletion at the *CSRNP2* locus resulted in loss of ZFP143 binding at the *CSRNP2* promoter and an ∼1.7-fold reduction in PRO-seq signal (**Fig. 3B**). To evaluate the hypothesis that ZFP143’s effects are restricted to P-P, E-P, or ZFP143-bound loops we classified loops based off H3K27ac^90^, H3K4me1^90^, and ZFP143^49^ ChIP-seq data in HEK293T cells and performed APA, again finding little to no change in loop strength upon ZNF143 depletion (**Fig. 3C**). Additionally, loop strengths were also largely unchanged between the untreated and ZNF143 depletion conditions, and insulation scores were unaffected by ZFP143 depletion (**Fig. 3D-E**).

**Figure 3.**
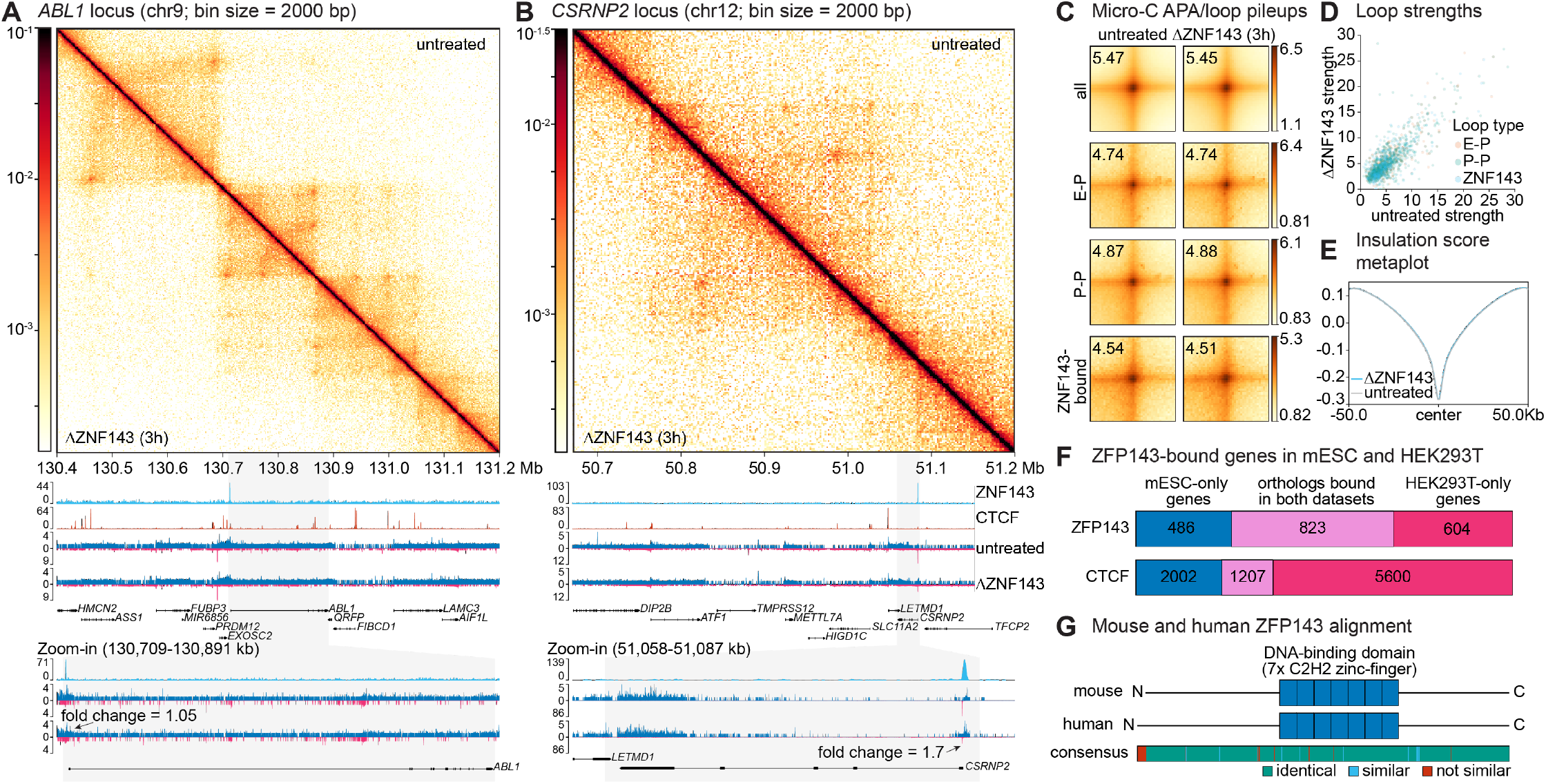
Loops are unaffected by ZNF143 depletion in HEK293T cells. (**a**-**b**) Panels showing genomics data at two representative loci in HEK293T clone 30. Shown from top to bottom are Micro-C maps at 2000 bp resolution comparing untreated to ΔZNF143 (3h), ZNP143 ChIP-seq tracks (cyan), CTCF ChIP-seq tracks (red), and PRO-seq with plus strand in blue and minus strand in magenta for untreated and ΔZNF143 (3h). Shown below, are zoom-ins on ZNF143-bound genes. (**a**) Shown is the *ABL1* locus. (**b**) Shown is the *CSRNP2* locus. (**c**) Aggregate peak analysis (APA)/loop pileup analysis for all called loops, E-P loops, P-P loops, and ZNF143-bound loops across untreated and ΔZNF143 (3h) conditions in log_10_ scale. Shown in the upper left corner is the average signal within the plot. (**d**) Scatterplots of boundary strengths calculated as in (**c**) for E-P loops (orange), P-P loops (teal), and ZNF143 loops (cyan) comparing ΔZNF143 (3h) to untreated. (**e**) Metaplots over insulation score tracks called at 50 kb resolution comparing untreated (gray) and ΔZNF143 (3h) (cyan). (**f**) Venn diagrams showing high levels of overlap in genes with ZFP143 or CTCF bound at their promoters between mESC and HEK293T ChIP-seq datasets. (**g**) Diagrams of mouse and human ZFP143 showing the location of the seven C2H2-zinc finger domains. Below the relationship between the two amino acid sequences is shown with identical amino acids shown in teal, similar amino acids shown in cyan, and dissimilar amino acids shown in red.

Given the conserved absence of loop function in both mouse and human cells, we asked if ZFP143’s binding exhibits evolutionary conservation in other ways. To this end, we took the set of genes bound by ZFP143 in mESCs and in HEK293T cells and identified ortholog pairs in both sets^91^. We found that 63% of genes bound by ZFP143 in mESCs are also bound in HEK293T cells despite methodological differences between the ChIP-seq datasets and the use of two very distinct cell types, a level of conservation at genes greater even than CTCF (**Fig. 3F**). Additionally, we found ∼97% sequence identity and ∼99% sequence similarity between the mouse and human ZFP143 protein sequences (**Fig. 3G**). Taken together, these results indicate that while there is no significant role for ZFP143 as a looping factor in either mESCs or HEK293T cells, ZFP143 binding is nevertheless heavily conserved.

### ZFP143 binds largely independently of CTCF

We next investigated the mechanism of ZFP143’s binding. It has been hypothesized that ZFP143 binds alongside CTCF or is recruited by CTCF to mediate looping^27,29^. To evaluate this hypothesis, we performed ChIP-seq against both ZFP143 and CTCF in all mESC clones under all depletion conditions, finding strong agreement between clones and conditions (**Fig. S4A-D**) as well as successful protein depletions (**Fig. S4E-G**). We first examined the overlap between called CTCF and ZFP143 ChIP-seq peaks in the untreated condition and found only 184 shared peaks representing ∼11.4% of all ZFP143 peaks and only ∼0.3% of all CTCF peaks (**Fig. 4A**). Due to the small overlap, we next asked if ZFP143 binding tends to overlap other features such as transcription start sites (TSSs) or enhancers we previously annotated. We found that 78.6% of ZFP143 peaks are within 1 kb upstream of an annotated TSS and 12.4% overlap an enhancer (**Fig. 4B**). Conversely, CTCF binding is largely inter-or intragenic. We next asked whether acute ZFP143 depletion affects CTCF binding or vice versa. Of the 60,498 CTCF peaks, none were significantly changed upon three-hour ZFP143 depletion, and of the 1,615 called ZFP143 peaks, 40 decreased in strength upon three-hour CTCF depletion (**Fig. 4C**). To assess whether the ZFP143 peaks changed in response to CTCF directly we computed the number of significantly decreased ZFP143 ChIP-seq peaks that overlapped a CTCF peak and found that 39 of the 40 decreased peaks indeed overlapped a CTCF peak. To confirm that ZFP143 or CTCF depletion does not affect the binding profile of CTCF or ZFP143 apart from peak height, we generated metaplots over ChIP-seq peaks in both datasets (**Fig. 4D**). We again found a small decrease in ZFP143 signal upon CTCF depletion only for those loci bound by CTCF and no change in CTCF ChIP-seq upon ZFP143 depletion with no change in the shape of the peaks.

**Figure 4.**
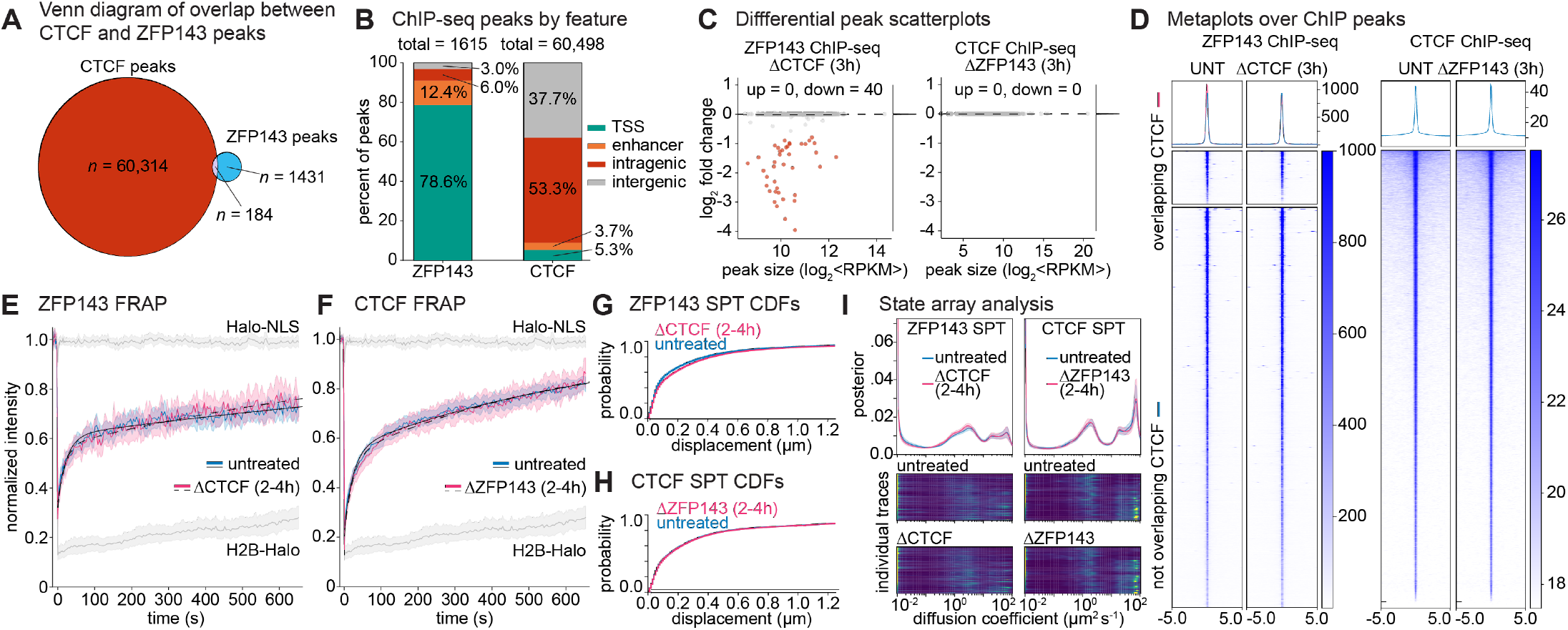
ZFP143 largely binds independently to CTCF. (**A**) Venn diagrams of the overlap in CTCF and ZFP143 binding between HEK293T and mESCs assessed by ChIP-seq. (**B**) Stacked bar chart showing the percentage of ZFP143 and CTCF ChIP-seq peaks categorized as overlapping TSSs (teal), enhancers (orange), intragenic (red), or intergenic (gray). (**C**) Scatterplots showing the log_2_ fold change of read counts at ChIP-seq peaks between either ΔCTCF (3h) and untreated ZFP143 ChIP-seq (left) or ΔZFP143 (3h) and untreated CTCF ChIP-seq (right) as a function of peak size. The number of significantly increased or decreased peaks as determined by DESeq2 (*p*_adj_ ≤ 0.05 and FC ≥ 1.5) is given at the top, and significantly changed peaks are colored in red. A histogram showing the density of points is given on the right of each plot. (**D**) Metaplots over ChIP-seq peaks for untreated or ΔCTCF (3h) ZFP143 ChIP-seq (left) or untreated or ΔZFP143 (3h) CTCF ChIP-seq (right). In the left plot, peaks are split into those overlapping a CTCF peak (top panel) for which the metaplot is shown in magenta and those not overlapping a CTCF peak (bottom panel) for which the metaplot is shown in blue. (**E-F**) FRAP curves showing normalized intensity as a function of time, and Halo-NLS and H2B-Halo controls are shown in gray. Error bars indicate 95% confidence intervals. (**E**) FRAP curves of ZFP143 in the untreated (blue) condition versus ΔCTCF (2 to 4 hours; magenta) averaging clones A and B. (**F**) FRAP curves of CTCF in the untreated (blue) condition versus ΔZFP143 (2 to 4 hours; magenta). (**G-H**) CDFs of jump displacement probabilities from SPT shown for a 24 ms time-lag. (**G**) SPT of ZFP143 comparing ΔCTCF (2 to 4 hours; magenta) with untreated (blue). (**H**) SPT of CTCF comparing ΔZFP143 (2 to 4 hours; magenta) with untreated (blue). (**I**) State-array SPT (saSPT) output showing the posterior distribution as a function of diffusion coefficient in µm^2^ s^-1^ for ZFP143 SPT (left) and CTCF SPT (right). Curves give the mean across traces and the shaded area gives the 95% percentile across all movies analyzed. Heatmaps show the individual traces averaged in the curve for all movies analyzed. Either ΔCTCF (2 to 4 hours) or ΔZFP143 (2 to 4 hours) is shown in magenta, and untreated is shown in blue.

Having established that ZFP143 and CTCF largely do not influence one another’s binding loci by ChIP-seq, we sought to test the hypothesis that they influence each other in terms of residence time or bound fraction. Here, we again performed FRAP on ZFP143 and CTCF upon a two-to-four-hour depletion of CTCF or ZFP143 respectively. We found that depleting CTCF does not appreciably change ZFP143’s FRAP recovery and depleting ZFP143 also does not change CTCF’s recovery (**Fig. 4E-F**). Next, to evaluate the influence of CTCF depletion on ZFP143’s bound fraction, we performed SPT of ZFP143 upon a two-to-four-hour depletion of CTCF finding a similar jump displacement distribution and a small decrease in mean bound fraction of ∼5 percentage points (**Fig. 4G**). SPT of CTCF under ZFP143 depletion conditions similarly revealed little change (**Fig. 4H**). To further test this result, we generated distributions of diffusion coefficients across all movies using state array SPT (saSPT), a method that uses variational Bayesian inference to extract the most likely diffusing states underlying a dataset^92^. Here, we again found strong agreement between diffusion rates in ZFP143 or CTCF SPT with or without CTCF or ZFP143, respectively (**Fig. 4I**). Overall, ZFP143 binding appears to have little effect on CTCF binding. However, while the effect of CTCF binding on ZFP143 is small, the loss of ZFP143 signal at ∼2.5% of sites upon CTCF depletion is consistent with our observed reduction in bound fraction. We conclude that ZFP143 and CTCF largely bind chromatin independently, except in some cases where they have binding motifs near each other.

### ZFP143 is a transcriptional regulator of ribosomal genes

Due to the observation that ZFP143 is highly conserved and binds preferentially to the promoter regions of genes, we next asked if and how ZFP143 functions as a transcriptional regulator consistent with prior work^46–53^. To understand ZFP143’s binding relative to TSSs, we called most-likely motifs for each ZFP143 ChIP-seq peak and plotted distributions of these motifs relative to annotated TSSs for peaks overlapping the 2 kb surrounding the TSS (**Fig. 5A**). We found that over 70% of motifs are upstream of the promoter with a median distance of 38 bp between the TSS and the center of the ZFP143 motif. Since ZFP143 binding is heavily enriched for sites nearby TSSs, we assessed changes to transcription using PRO-seq, with depletion of ZFP143, CTCF, and both ZFP143 and CTCF for 3 hours and found strong agreement between related clones and replicates (**Fig. S5A-C**). We first asked how nascent RNA transcripts are distributed around ZFP143 binding sites. Here, we generated metaplots of PRO-seq signal centered on called ZFP143 motifs as above and found a peak in signal in front of and behind the ZFP143 motif with a preference for transcripts on the same strand as the called motif (**Fig. 5B, Fig. S5D**). Additionally, we found that ZFP143 motifs upstream of the TSS exhibit a ∼1.7-fold enrichment for being oriented on the same strand as the gene to which they are bound whereas no such strand bias exists for those binding downstream of the TSS (**Fig. 5C**). Overall, these data indicate a distance and strand-dependent mechanism for ZFP143’s transcriptional regulation.

**Figure 5.**
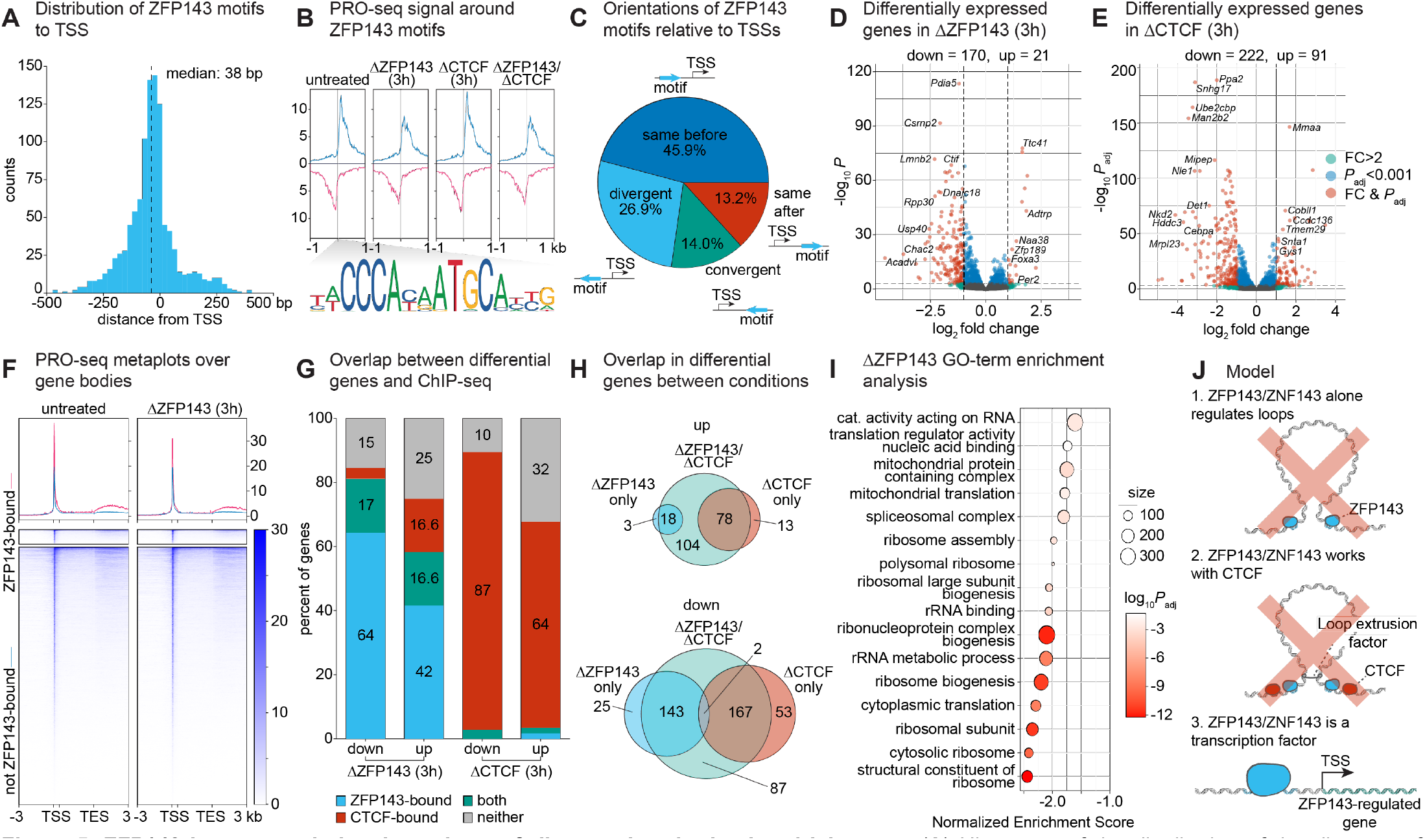
ZFP143 is a transcriptional regulator of ribosomal and mitochondrial genes. (**A**) Histogram of the distribution of the distance of ZFP143 motifs in ChIP-seq peaks overlapping + or – kb to a transcription start site (TSS) showing a median distance of 38 base pairs. (**B**) Metaplots over PRO-seq signal centered on ZFP143 motifs. Blue curves average the PRO-seq signal in the same strand as the given ZFP143 motif, and magenta curves average the PRO-seq signal on the opposite strand as the given ZFP143 motif. (**C**) Pie chart of the orientations of the ZFP143 motifs in (**A**) relative to their nearest TSS. Diagrams depict the ZFP143 motif relative to the TSS. (**D-E**) Volcano plots showing -log_10_ of the p adjusted value versus the log_2_ fold change of read counts of PRO-seq signal at annotated genes. Genes with a fold change greater than a factor of 2 are shown in green, genes with a p adjusted value less than 0.001 are shown in blue, and genes significant in both metrics are shown in red. The number of significantly changed genes evaluated by DESeq2 is shown above the plot. (**D**) Volcano plot showing the ΔZFP143 (3h) condition. (**E**) Volcano plot showing the ΔCTCF condition. (**F**) Metaplots over gene bodies showing average PRO-seq signal in the untreated (left) vs ΔZFP143 (3h; right) conditions. Genes with promoters bound by ZFP143 in ChIP-seq are shown in the upper panel and colored magenta in the metaplot, while non-ZFP143-bound genes are in the bottom panel and colored blue. Annotated TSSs and gene bodies are scaled to a fixed size for display, and data is shown plus or minus 3 kb from the transcription end site (TES) or TSS respectively. (**G**) Stacked bar plots of the percent of genes bound by either ZFP143 (cyan), CTCF (red), both (teal), or neither (gray) for the genes significantly changing expression in PRO-Seq as evaluated by DESeq2. (**H**) Venn diagrams of the overlaps in sets of genes significantly changing in the ΔZFP143 (3h; cyan), ΔCTCF (3h; red), or ΔZFP143/ΔCTCF (3h; teal) conditions. (**I**) A plot of significantly changing gene sets as a function of normalized enrichment score for GO-term enrichment analysis performed on the significantly changing genes in the ΔZFP143 (3h) condition. The size of the set is given by the size of the circle, and the points are colored by their log_10_ p adjusted value. (**J**) Possible models for ZFP143 function. It is unlikely that ZFP143 regulates loops alone (1) or with CTCF (2), but rather ZFP143 acts as a transcriptional regulator.

We next asked if acute perturbation of ZFP143 has direct transcriptional consequences. First, we compared the PRO-seq signal averaged over each gene between depletion conditions and the untreated condition. We found that acute ZFP143 depletion resulted in 170 significantly decreased genes and 21 significantly increased genes with a fold-change cutoff of two and an adjusted *p*-value cutoff of 0.001, a result that is similar in clone D, suggesting ZFP143 largely functions as a positive regulator of transcription (**Fig. 5D, Fig. S5E**). Similarly, acute depletion of CTCF resulted in 222 significantly decreased genes and 91 significantly increased genes, and depletion of both ZFP143 and CTCF exhibited no synergistic effects (**Fig. 5E, Fig. S5A, F**). Subsequently, we assessed the correspondence between ZFP143 binding and PRO-seq signal. To this end, we plotted metaplots over gene bodies of PRO-seq signal before and after three-hour ZFP143 depletion, and we found a reduction in PRO-seq signal for ZFP143-bound genes while other genes remain constant on average (**Fig. 5F**). We also asked what proportion of significantly increased or decreased genes upon three-hour ZFP143 or CTCF depletion have ZFP143 or CTCF bound within the 2 kb surrounding their TSS, finding that ∼81% of significantly decreased ZFP143 genes have ZFP143 bound compared to ∼59% for increased genes, again indicating that ZFP143 most often functions as a positive transcriptional regulator. Similarly, almost 90% of significantly decreased genes after CTCF depletion have CTCF bound compared to ∼65% for increased genes (**Fig. 5G**). Next, we hypothesized that since ZFP143 and CTCF largely bind independently to one another, they likely regulate different subsets of genes. Indeed, only two of the differentially expressed genes upon ZFP143 depletion are also differentially expressed upon CTCF depletion, and depleting both proteins largely reproduces the same sets as the two depletions do apart (**Fig. 5H**). Lastly, we asked what genes tended to be differentially expressed upon loss of ZFP143. We performed gene ontology enrichment analysis on the ΔZFP143 differentially expressed genes and found a strong enrichment for terms related to ribosomal function and mitochondrial proteins (**Fig. 5I**). Taken together, our results are consistent with a role for ZFP143 as a specialized transcription factor that regulates a few hundred genes often with ribosomal and mitochondrial function but otherwise functioning independently of CTCF and playing no major role in chromatin looping regulation.

## DISCUSSION

Here we show that ZFP143 is not a looping regulator in either mESCs or HEK293T cells as previously thought^17,25–40^, but rather it is a transcriptional regulator^41–53^. We find that ZFP143 anchors relatively few loops and those that it does are invariant to its acute depletion. Moreover, ZFP143 and CTCF overwhelmingly work independently from one another, and depletion of ZFP143 has no impact on CTCF ChIP-seq or its binding dynamics. Instead, we find that ZFP143 is highly conserved across species at the protein sequence level and in ChIP-seq binding in accordance with previous findings^31,42^, and it almost exclusively binds TSSs. In fact, ZFP143 preferentially binds promoters with a median distance of 38 bp to the nearest TSS and with the longest TF residence time yet measured to our knowledge. Additionally, acute depletion of ZFP143 causes primarily downregulation of a modest subset of genes enriched for ribosomal and mitochondrial function. Overall, our results are consistent with a model in which ZFP143 has no looping function either on its own or with CTCF; instead, ZFP143 binds promoters to regulate gene expression of an essential subset of genes (**Fig. 5J**).

ZFP143’s high level of evolutionary conservation and its tendency to bind promoters very stably near TSSs suggest that it could directly recruit the transcriptional machinery, a result consistent with prior work indicating that ZFP143 promoter binding is sufficient to drive gene expression in reporter gene assays^44–46,51^. Additionally, ZFP143 binding appears to be highly similar across tissues and cell types^93–96^, and it regulates expression of ribosomal and mitochondrial genes, which agrees with our finding that ZFP143 is essential in mESCs. Together, this result indicates that ZFP143 acts as an essential regulator of housekeeping genes without any looping function. Interestingly, this is also consistent with reports that housekeeping genes are generally not regulated by distal enhancer-promoter looping interactions^97,98^.

Gene expression in mammals is regulated by cell-type specific E-P interactions^1–8^. Therefore, ZFP143’s enrichment at promoters and its reported binding at loop/TAD boundaries alongside CTCF had offered an attractive explanation for how E-P interactions could be regulated: ZFP143 acts as an E-P/P-P-specific looping regulator possibly working with CTCF^17,25–30,32,34–38^. However, methodological limitations of prior work, which we overcome here, have impeded the field from completely understanding ZFP143’s function regulating genome architecture. In particular, acute depletion of ZFP143 allowed us to disentangle the direct effects of near-complete ZFP143 and/or CTCF loss from indirect effects, as compared to deletion or siRNA-mediated knockdown^76,77,99^. By using a double-degron approach, we were also able to combinatorially assay effects of ZFP143 on CTCF or *vice versa*. Moreover, we used Micro-C, a method that is both normalizable and uniquely sensitive to functional E-P and P-P interactions^32,73,74^, to accurately assay changes in looping interactions controlling for clone-to-clone and cell-type-to-cell-type variability. Consequently, though it cannot be ruled out that ZFP143 could regulate specific loops in some cell types or contexts, our results rule out ZFP143 as a *general* looping factor akin to CTCF. How then was ZFP143 misidentified as a general looping factor? ZFP143’s looping function and interaction with CTCF was predicated on ChIP-seq datasets using a single antibody from the ENCODE consortium^34^. However, as shown by Magnitov and Maresca *et al*. in their companion paper, this antibody is largely inconsistent with datasets using other antibodies including those using endogenously epitope tagged ZFP143 due to an apparent cross-reactivity with CTCF^100^. Thus, ZFP143 appears to have been originally misidentified as a looping factor mostly due to a single non-specific antibody.

ZFP143 was arguably the single most promising general looping factor candidate after CTCF and cohesin; however, it is not a general looping regulator. As a result, the picture of 3D genome structure regulation in mice and humans is on the one hand clear: CTCF acts as a general boundary factor organizing the genome into loops and domains^101^, something no other mouse or human protein likely does to a similar extent. On the other hand, since CTCF is expressed in all cell types, the question remains how cell-type specific domains and loops are established and regulated by protein factors. Our study helps to distinguish two broad models for E-P and P-P loop formation. One hypothesis is that general E-P looping factors exist which hold together E-P loops in most cases, similar to how CTCF forms most structural loops across cell types. However, most E-P loops are robust to loss of CTCF, as CTCF loss only affects the small subset of E-P and P-P interactions that have CTCF binding at either end^23^, Additionally, though other proteins have been suggested to perform this function to some extent^21,98,102–104^, these proteins do not fully explain the observed breadth of E-P and P-P contacts. Another possibility is that no single general E-P looping factor exists. Instead, E-P and P-P interactions may be mediated by the combined, multi-valent interactions among all the TFs, co-activators, and other transcriptional proteins bound at promoter and enhancer regions^87^. This model explains both the absence of general E-P looping factors and how cell-type specific E-P and P-P loops result from cell-type specific TF expression. We therefore propose that, whereas structural CTCF/cohesin loops are mediated by two factors with very strong effects, E-P and P-P loops may instead form through a “strength-in-numbers” mechanism that integrates the combined affinity-based interactions of many weakly contributing transcriptional proteins.

## Acknowledgements

We thank Michael Guertin and Sathyan Kizhakke Mattada for generously sharing the previously generated HEK293T ZNF143-degron cell line^53^. We thank Elzo de Wit, Michela Maresca, and Mikhail Magnitov for coordinating publication and for helpful discussions. We thank Mustafa Mir for helpful discussion with microscope construction and Michael Davis for helpful discussion and his efforts with microscope construction. We thank Miles Huseyin for guidance on the Micro-C and ChIP-seq experiments and the genomics analysis as well as many helpful discussions and feedback on the manuscript. We thank Luke Lavis for providing Janelia Fluor dyes. We also thank Maxine Jonas, Jamie Drayton, Sumin Kim, Clarice Hong, Sarah Nemsick, and Asmita Jha for comments on the manuscript and the Hansen lab for discussions. We thank the MIT Koch Institute’s Robert A. Swanson (1969) Biotechnology Center for technical support, specifically the Integrated Genomics and Bioinformatics Core, flow core facilities, preclinical modeling facility, and the MIT BioMicroCenter, and this work was supported in part by the Koch Institute Support (core) Grant P30-CA014051 from the National Cancer Institute. We also thank the Walk-Up Sequencing services of the Broad Institute of MIT and Harvard. This work was primarily supported by NSF grant 2036037 and we acknowledge additional funding support from NIH grants DP2GM140938, R33CA257878, and UM1HG011536, NSF CAREER 2337728, Pew-Stewart Cancer Research Scholar grant, the Mathers Foundation, and the Gene Regulation Observatory at the Broad Institute. The raw data can be found at NCBI GEO under accession number GSE256246.

## Author contributions

D.N.N. and A.S.H. designed the project. D.N.N. built the single-molecule tracking microscope and conducted all experiments and analyses. A.S.H. supervised the project. Both authors contributed to drafting and editing the paper and figures.

