## Supplementary Information for "Putative Looping Factor ZNF143/ZFP143 is an Essential Transcriptional Regulator with No Looping Function"


**Contents**


- Supplementary Figures and Figure Legends
- Supplementary Table
- Materials and Methods
- References

### Supplementary Figures and Figure Legends

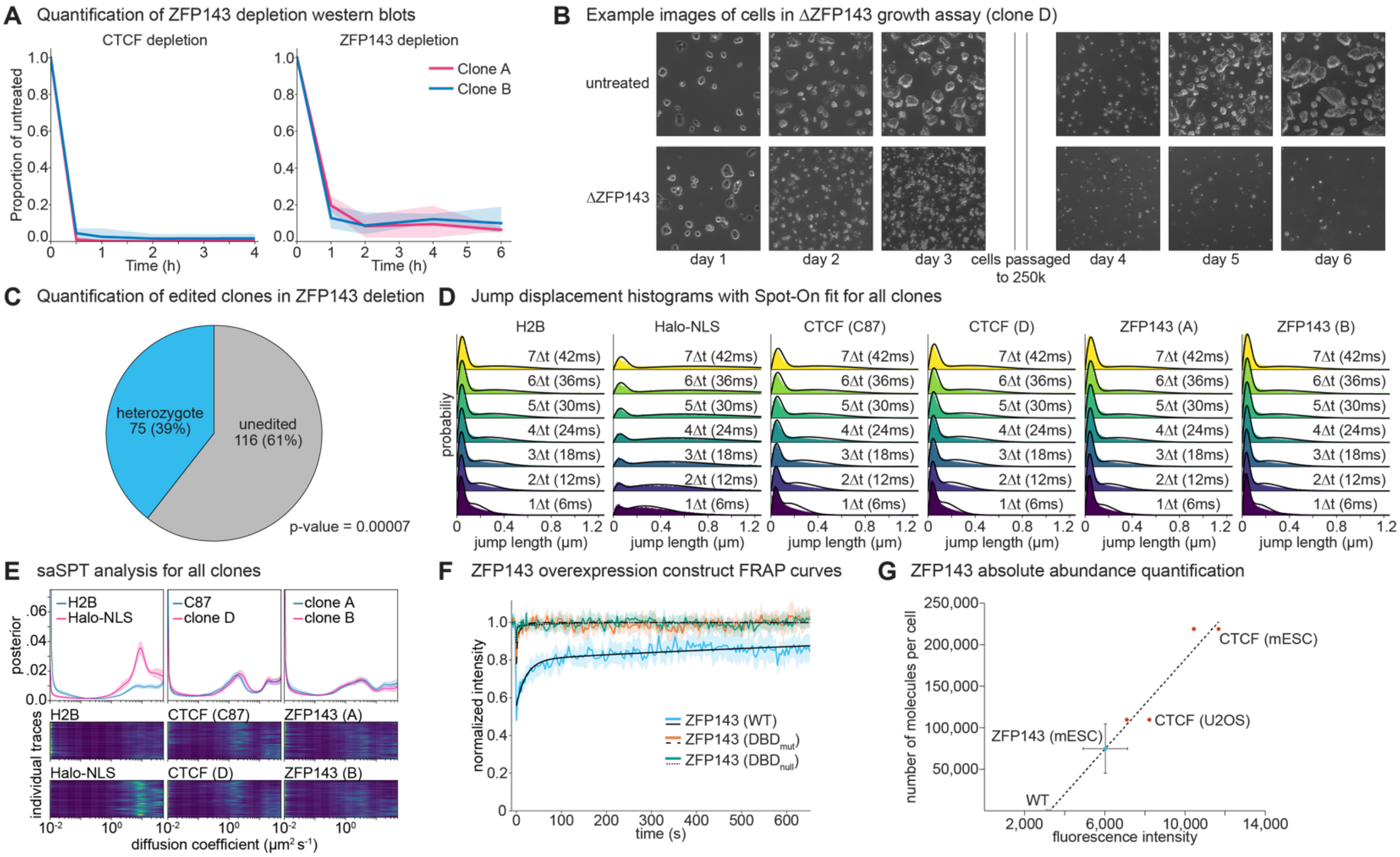


**Figure S1. ZFP143 and CTCF depletion is reproducible, and ZFP143 cannot be deleted.** (**A**) Quantification of depletion western blots showing the average loading control-normalized value for both CTCF and ZFP143 in clones A (magenta) and B (blue). Error bars indicate 95% confidence intervals. (**B**) Example images of mESCs during the growth assay performed in **Fig. 1D**. (**C**) A pie chart showing the number of clones that are homozygotes, heterozygotes, or unedited in 191 screened colonies for a ZFP143 deletion (p = 0.00007). The p-value was computed as the probability of getting no homozygotes assuming an equal probability of editing at each allele. (**D**) Jump displacement histograms with Spot-On fits for all experimental conditions assayed (H2B-Halo, Halo-3xNLS, C87 CTCF, clone D CTCF, clone A ZFP143, and clone B ZFP143). (**E**) saSPT plots showing the distribution of diffusion coefficients for the H2B-Halo and Halo-3xNLS binding controls (left), C87 and clone D CTCF (middle), and clones A and B ZFP143 (right). (**F**) FRAP of overexpressed, transfected ZFP143 binding constructs with representative FRAP fits. Error bars indicate 95% confidence intervals. (**G**) Plot showing the calibration curve used to calculate the ZFP143 absolute abundance as well as the quantification of ZFP143 absolute abundance. Error bars indicate standard deviation.

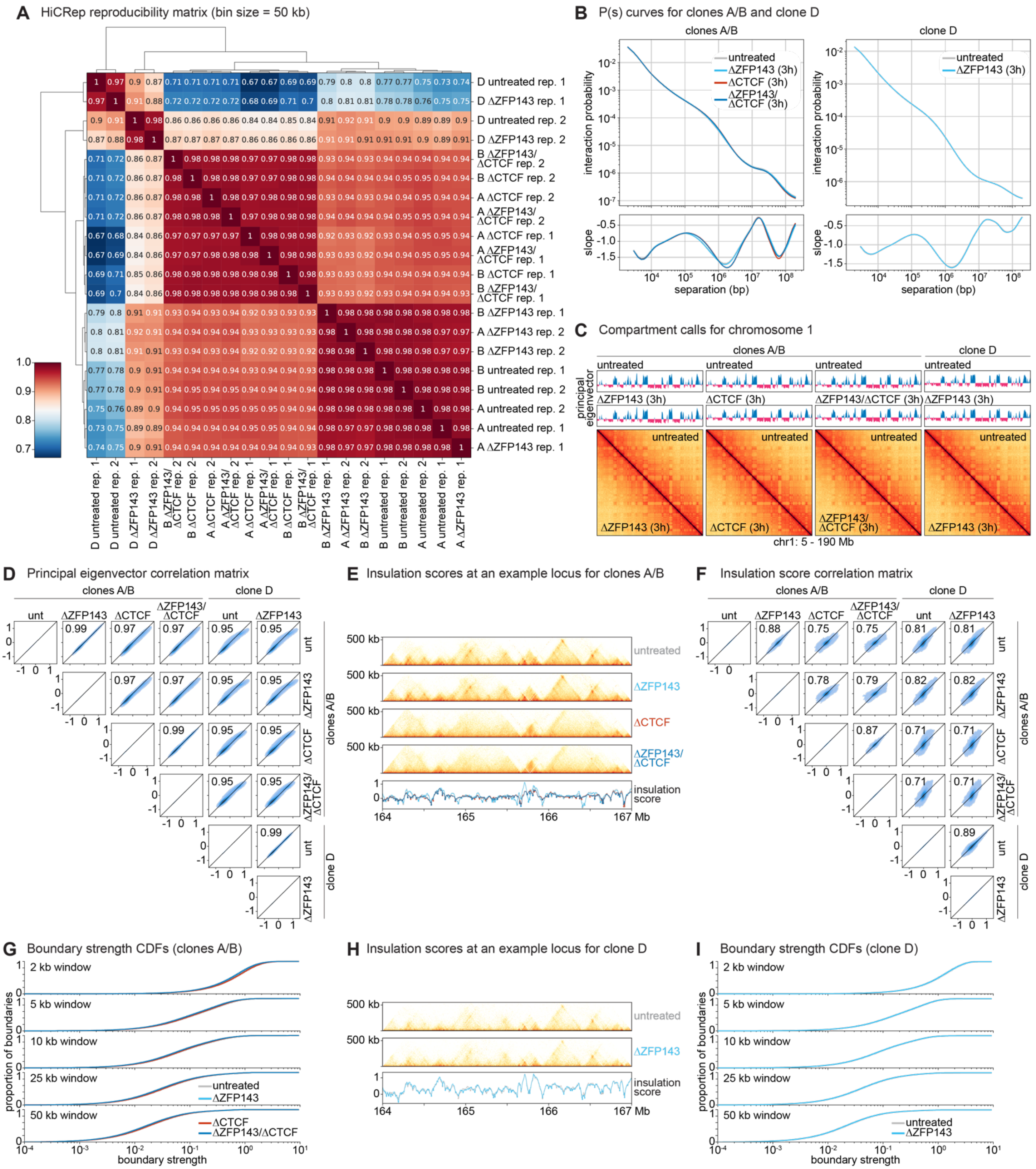


**Figure S2. Micro-C between mESC clones is highly reproducible.** (**A**) HiCRep reproducibility matrix for all mESC Micro-C conditions computed with 50 kb bin size. The values indicate stratum adjusted correlation coefficients between mESC conditions, and the black lines (left, top) show a hierarchical clustering based on the HiCRep values. (**B**) P(*s*) curves showing interaction probabilities at various scales for clones A and B (left) or clone (D) right for untreated (gray), ΔZFP143 3h (cyan), ΔCTCF 3h (red), or ΔZFP143/ΔCTCF 3h (blue) conditions. The bottom panel shows the derivative of the above plot. (**C**) Compartment calls for chr1:5-190Mb. The top panel shows principal eigenvector values for the below plots. Plots are shown at 500 kb resolution in $\log_{10}$ scale. (**D**) A matrix of Pearson correlations in principal eigenvector values between all clones. The blue shows a kernel density estimate for the data. (**E**) Insulation score tracks at a representative locus for clones A and B. The top four panels show the Micro-C at 5 kb resolution in $\log_{10}$ scale while the bottom panel shows insulation scores called at 50 kb resolution. (**F**) A matrix of correlations between insulation scores called at 50 kb resolution as in panel (**D**). (**G**) Cumulative distribution functions for insulation scores at boundaries, defined as minima in the insulation score, for insulation scores called at various resolutions in clones A and B. (**H**) Insulation score tracks at a representative locus for clone D. The top two panels show the Micro-C at 5 kb resolution in $\log_{10}$ scale while the bottom panel shows insulation scores called at 50 kb resolution. (**I**) Cumulative distribution functions for insulation scores at boundaries, defined as minima in the insulation score, for insulation scores called at various resolutions for clone D.

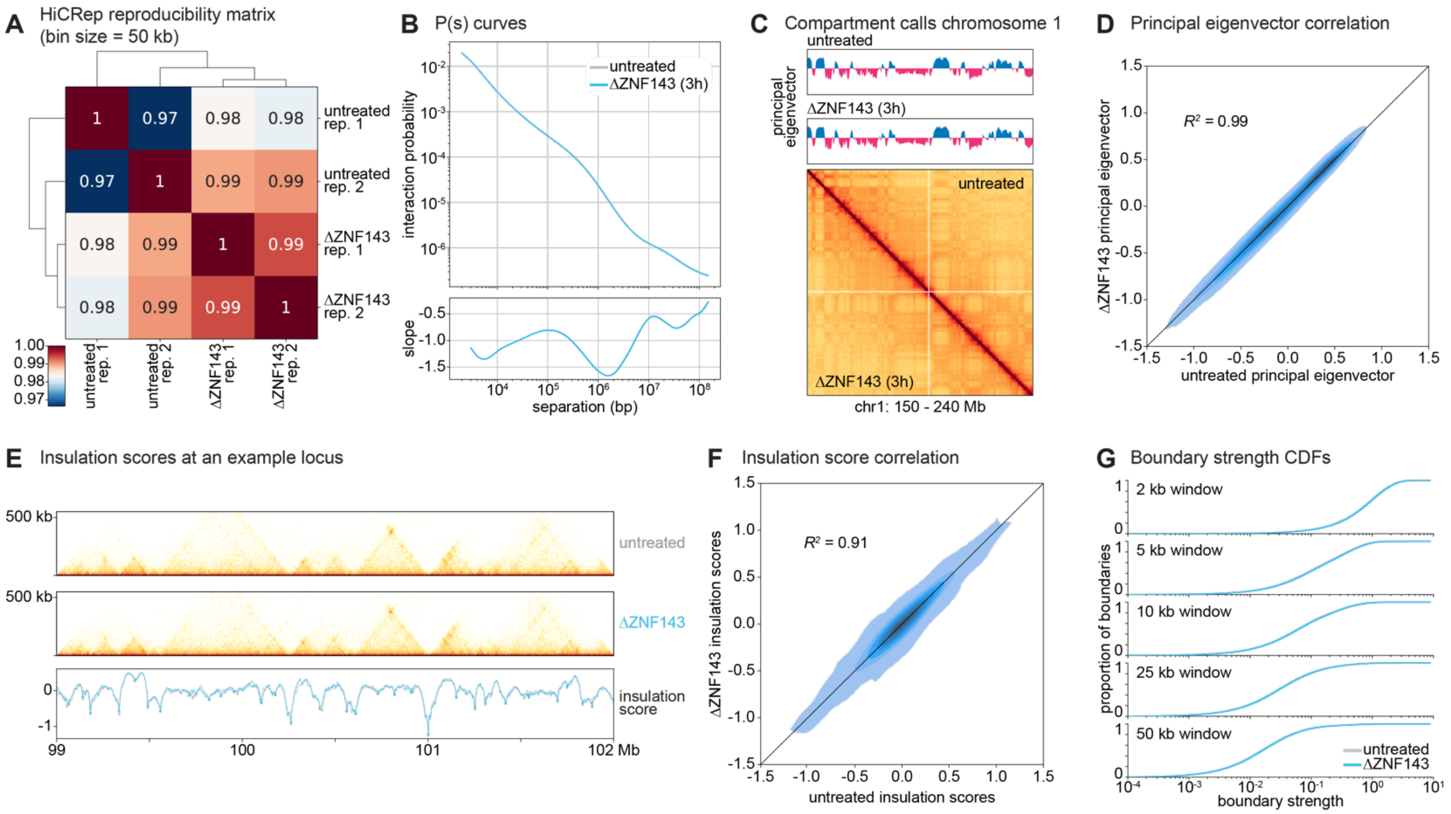


**Figure S3. Micro-C in the HEK293T clone 30 is highly reproducible.** (**A**) HiCRep reproducibility matrix for all HEK293T Micro-C conditions. The values indicate correlation coefficients between conditions, and the black lines (left, top) show a hierarchical clustering based on the HiCRep values. (**B**) P(*s*) curve showing interaction probabilities at various scales for clones 30 for untreated (gray) or ΔZFP143 3h (cyan) conditions. The bottom panel shows the derivative of the above plot. (**C**) Compartment calls for chr1:150-240Mb. The top panel shows principal eigenvector values for the below plots. Plots are shown at 500 kb resolution. (**D**) A kernel density estimate plot showing principal eigenvector values between untreated and ΔZFP143 3h clone 30. Values indicate Pearson correlation. (**E**) Insulation scores at a representative locus for clone 30. The top two panels show the Micro-C at 5 kb resolution in $\log_{10}$ scale while the bottom panel shows insulation scores called at 50 kb resolution. (**F**) A kernel density estimate plot of insulation scores called at 50 kb resolution as in panel (**D**). (**G**) Cumulative distribution functions for insulation scores at boundaries defined as minima in the insulation score as a function of boundary strength for insulation scores called at various resolutions.

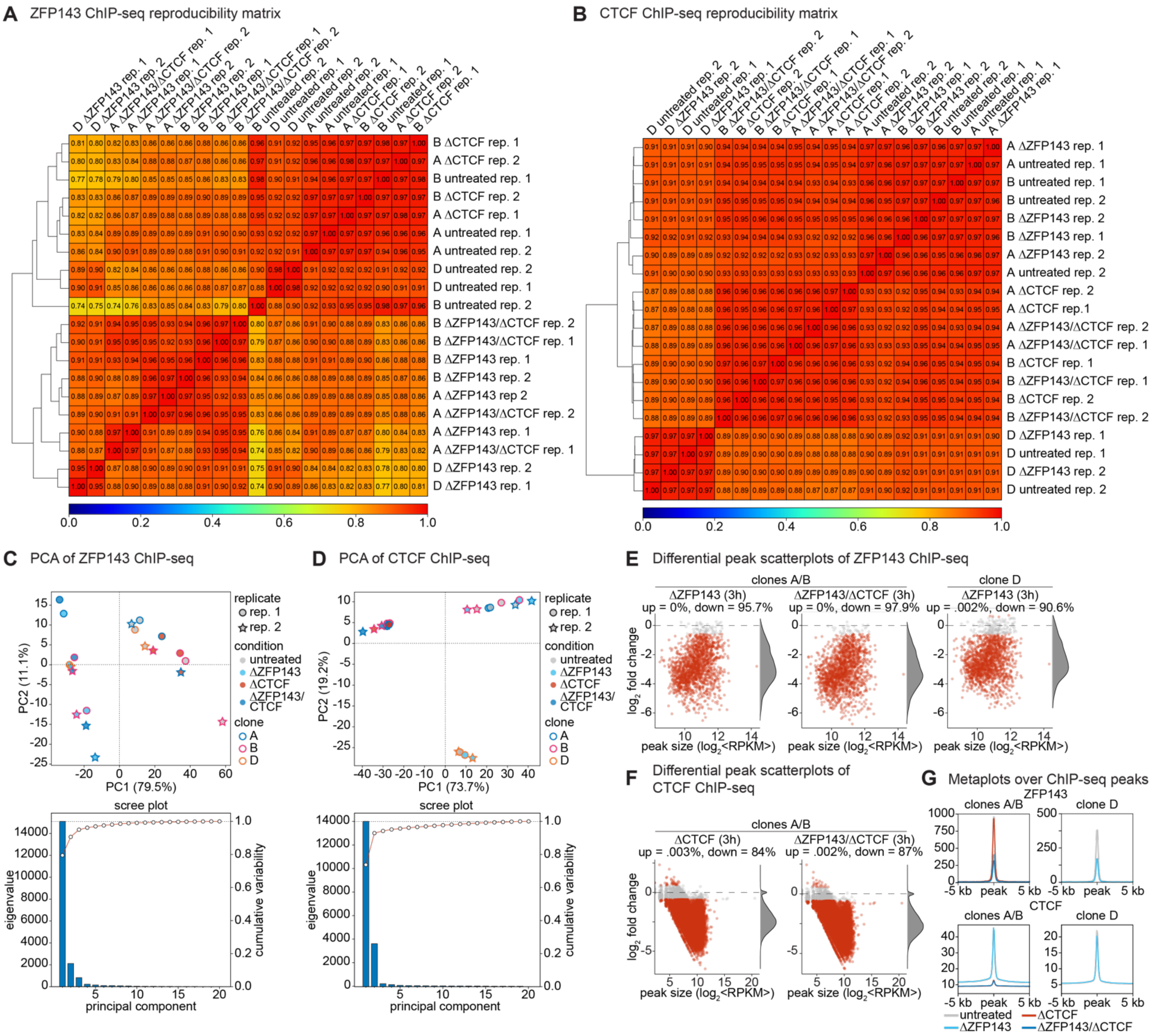


**Figure S4. ChIP-seq across all mESC conditions is highly reproducible.** (**A**-**B**) Reproducibility matrices between ChIP-seq conditions. The color and value indicate the correlation coefficient, and hierarchical clustering is shown on the left. (**A**) ZFP143 ChIP-seq. (**B**) CTCF ChIP-seq. (**C-D**) Principal component analysis (PCA) of ChIP-seq conditions. In the top plot, principal component 2 (PC2) is plotted against PC1 and the percent variance explained is shown in parentheses. The shape of the marker indicates replicate 1 (circle) or replicate 2 (star), while the color indicates untreated (gray), ΔZFP143 3h (cyan), ΔCTCF 3h (red), or ΔZFP143/ΔCTCF 3h (blue), and the outline color indicates clone A (blue), B (magenta), or D (orange). The bottom is a scree plot showing the eigenvalue of each principal component (blue bars, left axis) and the cumulative variability explained (red line, right axis). (**C**) ZFP143 ChIP-seq. (**D**) CTCF ChIP-seq. (**E-F**) Scatterplots showing the $\log_2$ fold change between the condition indicated at the top of the plot and the untreated condition as a function of peak size. Red dots indicate significantly changed peaks ($p_{adj} \leq 0.05$, fold change $\geq 1.5$). Histograms of point densities are shown to the right and the percent of peaks up or down is shown above the plot. (**E**) ZFP143 ChIP-seq. (**F**) CTCF ChIP-seq. (**G**) Metaplots over ChIP-seq peaks in clones A/B (left) or clone D (right) for ZFP143 ChIP-seq (top) or CTCF ChIP-seq (bottom) for untreated (gray), ΔZFP143 3h (cyan), ΔCTCF 3h (red), or ΔZFP143/ΔCTCF 3h (blue).

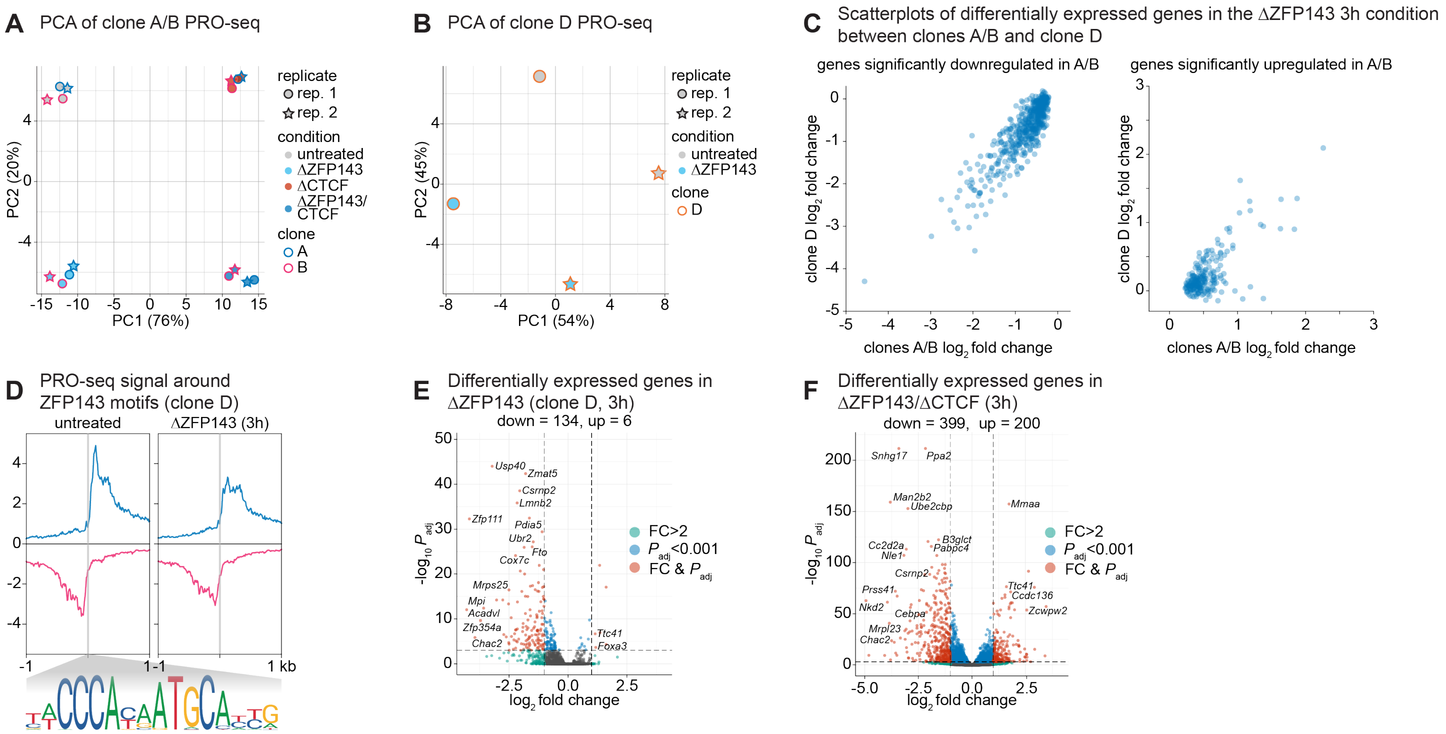


**Figure S5. PRO-seq across mESC conditions is highly reproducible, and the effects of ZFP143 and CTCF on gene expression are mainly additive.** (**A**-**B**) Principal component analysis (PCA) of PRO-seq conditions. Principal component 2 (PC2) is plotted against PC1 and the percent variance explained is shown in parentheses. The shape of the marker indicates replicate 1 (circle) or replicate 2 (star), while the color indicates untreated (gray), ΔZFP143 3h (cyan), ΔCTCF 3h (red), or ΔZFP143/ΔCTCF 3h (blue), and the outline color indicates clone A (blue), B (magenta), or D (orange). (**A**) PCA of PRO-seq for clones A and B. (**B**) PCA of PRO-seq for clone D. (**C**) Plots of the $\log_2$ fold change values between ΔZFP143 3h and untreated clone D and clones A/B for genes significantly downregulated in clones A/B (left) or significantly upregulated in clones A/B (right) upon 3-hour ZFP143 depletion showing strong correlation. (**D**) Metaplots over PRO-seq signal centered on ZFP143 motifs in clone D PRO-seq. The blue curves average the PRO-seq signal in the same strand as the given ZFP143 motif, and magenta curves average the PRO-seq signal on the opposite strand as the given ZFP143 motif. (**E-F**) Volcano plots showing $-\log_{10}$ of the p adjusted value versus the $\log_2$ fold change of read counts of PRO-seq signal at annotated genes. Genes with a fold change greater than a factor of 2 are shown in green, genes with a p adjusted value less than 0.001 are shown in blue, and genes significant in both metrics are shown in red. The number of significantly changed genes evaluated is shown above the plot. (**E**) Genes significantly changed upon 3-hour depletion of ZFP143 in clone D. (**F**) Genes significantly changed upon 3-hour depletion of both ZFP143 and CTCF in clones A/B.

#### Supplementary Tables

**Table S1. List of oligonucleotides and primers.**
**Table S2. List of Micro-C loops overlapping ZFP143 in mESCs.**
**Table S3. List of Micro-C loops in HEK293T cells.**
**Table S4. List of Micro-C E-P loops in HEK293T cells.**
**Table S5. List of Micro-C P-P loops in HEK293T cells.**
**Table S6. List of Micro-C loops overlapping ZFP143 in HEK293T cells.**
**Table S7. List of ZFP143 ChIP-seq peaks in mESCs.**
**Table S8. List of CTCF ChIP-seq peaks in mESCs.**
**Table S9. List of ZFP143 ChIP-seq peaks in HEK293T cells.**
**Table S10. List of CTCF ChIP-seq peaks in HEK293T cells.**
**Table S11. List of putative enhancers in mESCs.**
**Table S12. List of putative enhancers in HEK293T cells.**
**Table S13. DESeq2 output comparing ΔZFP143 to untreated PRO-seq in clones A and B.**
**Table S14. DESeq2 output comparing ΔCTCF to untreated PRO-seq in clones A and B.**
**Table S15. DESeq2 output comparing ΔZFP143/ΔCTCF to untreated PRO-seq in clones A and B.**
**Table S16. DESeq2 output comparing ΔZFP143 to untreated PRO-seq in clone D.**

The tables can be downloaded here:

**Supplementary Tables S1-S12:**
https://www.dropbox.com/scl/fi/wynzit62dot80hosvtv9x/Supplementary_Tables_1_to_12.xlsx?rlkey=h79hswhaiyvf97k2gnffuo8bf&dl=0

**Supplementary Tables S13-S16:**
https://www.dropbox.com/scl/fi/wp4l1poqyv4om0h8pykc8/Supplementary_Tables_13_to_16.xlsx?rlkey=c1id7ifoh8isy6gvzwt8ojjir&dl=0

### Materials and Methods

#### Data and code availability

- Micro-C, ChIP-seq, and PRO-seq data have been deposited at GEO and are publicly available at https://www.ncbi.nlm.nih.gov/geo/query/acc.cgi?acc=GSE256246
- Original code reported in this paper is available at the following Github link: https://github.com/ahansenlab/ZNF143_analysis_code

#### Cell culture

Mouse embryonic stem cells (JM8.N4 mESC[105]; RRID:CVCL_J962) obtained from the KOMP Repository at the University of California Davis were cultured under 5% CO2 at 37°C under feeder-free conditions. The mESCs were grown in medium consisting of KnockOut DMEM (ThermoFisher #10829-018) supplemented with 15% FBS (Avantor Seradigm #89510-186, Lot #190B20)1000 U mL-1 LIF (homemade)[12], 1 mM MEM Non-Essential Amino Acid Solution (ThermoFisher #11140050), 2 mM GlutaMAX™ supplement (ThermoFisher #35050061), 100 µg mL-1 Penicillin-Streptomycin (ThermoFisher #15140122), 0.1 mM 2-mercaptoethanol (ThermoFisher #31350010), 10 µM MEK inhibitor (Tocris #PD0325901), and 3 µM GSK inhibitor (Sigma-Aldrich #SML1046) on plates pre-coated with sterile 0.1% gelatin (Sigma-Aldrich, G1890). The mESCs were passaged every 2 days by dissociation with TrypLE Express Enzyme (ThermoFisher #12605028), and half the medium was replaced daily. Cells were tested regularly for the presence of mycoplasma at the MIT Swanson Biotechnology Center Preclinical Modeling Facility.

For all imaging experiments, cells were plated on 35 mm no. 1.5 glass-bottom imaging dishes (MatTek #P35G-1.5-14-C) pre-coated with 0.1% gelatin. Directly prior to imaging, the medium was changed to phenol red free DMEM with all other components of the medium the same. For transfection imaging conditions, cells were transfected using Lipofectamine 2000 (ThermoFisher #11668027) according to the manufacturer's protocol 16 to 24 hours prior to imaging.

The human cell lines C32[12] (U2OS), Clone 30[53] (HEK293T), and HEK293T cells (ATCC #CRL-3216) were cultured under 5% CO2 at 37°C in medium consisting of high glucose DMEM (ThermoFisher #11965092) supplemented with 10% FBS (Avantor Seradigm #89510-186, Lot #190B20) and 100 µg mL-1 Penicillin-Streptomycin (ThermoFisher #15140122). The cells were passaged every 2-4 days by dissociation with TrypLE Express Enzyme (ThermoFisher #12605028).

#### Protein depletion

For depletion of ZFP143 in the clone A and clone B cell lines, the cells were incubated in media containing 100 µM 5-Ph-IAA (MedChemExpress #HY-134653) and 300 nM HaloPROTAC3 (Promega # GA3110), and for depletion of CTCF in the same lines the, the cells were incubated in media containing 100 nM dTAG-13 (Tocris # 6605). For the depletion of ZFP143 in the clone D cell line, the cells were incubated in media containing 100 nM dTAG-13, and for the depletion of ZNF143 in the clone 30 HEK293T cell line, the cells were incubated in media containing 500 µM IAA.

#### CRISPR/Cas9-mediated genome editing

CRISPR/Cas9-mediated genome editing of mESCs was performed largely according to published protocols[12,106]. Briefly, cells were co-transfected in a 6-well dish with a Cas9 plasmid encoding a puromycin resistance gene and the sgRNA (Addgene #62988) and a repair construct plasmid comprised of the inserted DNA sequence flanked on each side by 500bp of homology to the target gene in a 1:3.5 µg ratio using Lipofectamine 2000 (ThermoFisher #11668027) according to the manufacturer's protocol. The sgRNAs were designed using the CRISPOR online tool[107] and are listed in **Table S1**. The day following transfection, cells were passaged to a new plate at single-cell densities with 1 µg $mL^{-1}$ puromycin. Following a 36h selection, puromycin was removed and cells were grown for approximately 1 week, after which individual clones were transferred to a 96-well plate and expanded.

The homozygous edited clones were identified by a PCR screen from genomic DNA (primers listed in **Table S1**) using a 3 primer PCR strategy (2 primers external to the homology arm and 1 internal to the inserted sequence). To prepare genomic DNA, cells were scraped and resuspended into buffer containing 10 mM Tris-HCl pH8, 1 mM EDTA, 25 mM NaCl, and 200 µg $mL^{-1}$ proteinase K (Viagen Biotech #501-PK) and incubated at 60°C for 1 hour followed by 95°C for 10 minutes. The edited clones were subsequently expanded and high-quality genomic DNA was obtained via phenol-chloroform extraction followed by ethanol precipitation. Approximately 1 million cells were lysed in 50 µL buffer containing 10 mM NaCl, 10 mM Tris pH7.5, 10 mM EDTA pH8.0, 0.5% N-lauryl sarcosine, and 1 mg $mL^{-1}$ proteinase K at 55°C for 4 hours. The lysate was mixed with an equal volume of phenol:chloroform:isoamyl alcohol (Sigma-Aldrich #P2069-100ML) and suspended in a phase lock tube (Quantabio #2302820). After centrifugation, the aqueous phase was recovered, and the gel was washed with an equal volume of chloroform. Upon addition of a 1:10 volume of 3M sodium acetate pH5.5 solution as well as 2 volume equivalents of 100% ice-cold ethanol, the DNA was incubated at –80°C for 30 minutes. The precipitated DNA was isolated by centrifugation and resuspended in deionized water. Homozygous editing was confirmed by PCR followed by sanger sequencing of the expected product. For protein-tagged cell lines, the edit was confirmed by western blot and imaging.

#### Growth assay

To estimate the growth rate of cells in the absence of ZFP143, clone D cells were counted using a Countess 3 automated cell counter (ThermoFisher #AMQAX2000) and seeded at a uniform density of 250,000 cells per well of a 6-well plate according to the culturing conditions above. To maintain continuous degradation of the protein, media was changed every day on all wells of the 6-well dish. Each day at the same time, the cells were counted and imaged using the EVOS cell imaging system (ThermoFisher #AMF5000), and after the third count cells were seeded at a uniform density of 250,000 cells per well of a 6-well plate as above. The assay was carried out until there were fewer than 250,000 cells remaining.

#### Western blotting

Protein extracts were prepared from nuclei isolated from cultured cells. To isolate nuclei, pelleted cells were suspended in 10 volumes of buffer containing 10 mM HEPES pH 7.9, 1.5 mM $MgCl_2$, 10 mM KCl, 0.5 mM dithiothreitol, 0.5 mM phenylmethylsulfonyl fluoride, and 1X cOmplete protease inhibitor (Roche #11873580001), incubated on ice for 10 minutes, and pelleted by centrifugation at 4°C and 1500g for 5 minutes. The pellet was suspended in 3 volumes of the above buffer supplemented with 0.1% IGEPAL CA-630 (Sigma-Aldrich #I8896-100ML) and pelleted by centrifugation as above. To prepare the protein extract, the pellet was suspended in 1 volume of buffer containing 5 mM HEPES pH 7.9, 26% glycerol, 1.5 mM $MgCl_2$, 0.2 mM EDTA, 0.5 mM dithiothreitol, 1X cOmplete protease inhibitor, and 250 mM NaCl. The NaCl concentration was adjusted to 400 mM by dropwise addition of 5 M NaCl and the solution was incubated at 4°C for 1 hour. The solution was pelleted by centrifugation at 4°C and 2500g for 20 minutes, and the supernatant was taken as the protein extract. Protein concentrations were determined using the Qubit protein assay kit (ThermoFisher #Q33212).

For western blotting, protein extracts totaling 15 µg were combined with loading buffer (2% SDS, 100 mM Tris pH 6.8, 100 mM dithiothreitol, 10% glycerol and 0.1% bromophenol blue) and boiled at 95°C for 10 minutes. The extracts were loaded on a 4–15% Mini-PROTEAN TGX precast protein gel (Bio-Rad #4561085) with the Precision Plus Protein dual color standards (Bio-Rad #1610374S) in 1X Tris/Glycine/SDS running buffer prepared according to manufacturer instructions (Bio-Rad #1610732). After running, the protein was transferred to a 0.2 µm nitrocellulose membrane (GenScript #L00732) using the eBlot L1 fast wet transfer system on the standard setting (GenScript #L00686).

After the transfer, membranes were blocked by 30-minute incubation in 5% nonfat dry milk (Genesee Scientific #20-241) in TBS-T (TBS with 0.1% Tween 20 (VWR #M147-1L)). The membranes were then incubated overnight at 4°C with the following antibodies suspended in TBS-T with 5% milk: CTCF 1:1000 (Millipore #07-729), ZNF143 1:1000 (Novus Biologicals H00007702-M01), TBP 1:3000 (Abcam #1TBP18). The secondary incubation was performed at room temperature for 1.25 hours by incubation with the following antibodies suspended in TBS-T with 5% milk: Mouse IgG HRP linked 1:5000 (Cytiva #NA931-1ML) or Rabbit IgG HRP linked 1:5000 (Cytiva #NA934-1ML). The blots were imaged by chemiluminescence with Pierce ECL western blotting substrate (ThermoFisher #32209) and measured with the ChemiDoc MP imaging system. Band intensity quantification was performed in Image Studio Lite.

#### Micro-C library preparation

To assay the change in looping upon loss of ZFP143/ZNF143 and CTCF, a 3-hour depletion of the proteins was performed as described above. The micro-C library preparation was performed according to a previously published protocol[87]. Briefly, 5 million cells were crosslinked for 45 minutes with 3 mM disuccinimidyl glutarate at room temperature. In the final 10 minutes of crosslinking, formaldehyde was added to a final concentration of 1%. After 45 minutes, the reaction was quenched with 0.375 M Tris pH 7.5, and the pellets were snap frozen. The crosslinked chromatin was digested to mononucleosomes using micrococcal nuclease (Worthington Biochemical # LS004798) at a concentration determined by titration. Biotin-dATP and biotin-dCTP were incorporated into the digested chromatin ends in a reaction with 1 U per 1 μg DNA of Klenow fragment (New England Biolabs #M0210L). The chromatin ends were proximity ligated in a reaction with 20 U $\mu l^{-1}$ T4 DNA ligase (New England Biolabs #M0202M) and the biotin-dNTPs were removed from the unligated ends using 5 U $\mu l^{-1}$ Exonuclease III (New England Biolabs #M0206L). Finally, the crosslinking was reversed by incubation overnight at 65°C in a solution containing 2 mg $mL^{-1}$ proteinase K (Viagen Biotech #501-PK), 1% SDS, 250 mM NaCl, and 0.1 mg $mL^{-1}$ RNase A (ThermoFisher # EN0531).

The DNA was subsequently purified using the Zymo DNA Clean and Concentrator (DCC-25) kit (Zymo Research #D4034). The purified DNA was run on a 1% agarose gel and the di-nucleosomal band was size selected by cutting the band between 250 and 400bp. The DNA from the band was purified using the Zymoclean Gel DNA recovery kit (Zymo Research #D4008), and proximity ligated DNA was isolated by streptavidin bead purification with Dynabeads MyOne Streptavidin T1 (ThermoFisher #65601). The purified DNA proceeded to library preparation following the NEBNext Ultra II Library Prep kit (New England Biolabs #E7645L), and the libraries were purified using 0.9X Ampure XP bead purification (Beckman Coulter #A63881). Library quality was assessed via Fragment Analyzer and qPCR at the MIT BioMicro Center. Biological duplicates were performed for all Micro-C libraries, and libraries were pooled in a 1:1 molar ratio based on the qPCR results and sequenced on either a NovaSeq 6000 S1 flow cell or a NovaSeq X flow cell at the Broad Institute of MIT and Harvard.

#### Chromatin Immunoprecipitation Sequencing (ChIP-seq) library preparation

To assay the effects of acute ZFP143/ZNF143 and CTCF depletion on CTCF or ZFP143/ZNF143 binding, a 3-hour depletion of the proteins was performed as described above. The ChIP-seq experiment was performed as previously described[108]. First, chromatin was isolated from crosslinked tissue culture cells. 50 million cells were dissociated, pelleted, and suspended at a concentration of 5 million cells per mL in PBS. Formaldehyde was added to a final concentration of 1%, and the cells were incubated at room temperature for 15 minutes with rotation. The reaction was quenched with the addition glycine at a concentration of 125 mM at room temperature for 5 minutes. The cells were snap-frozen in liquid nitrogen.

Subsequently nuclei were isolated. The crosslinked cells were first suspended in 10 mL buffer containing 10% glycerol, 50 mM HEPES-KOH pH 7.9, 140 mM NaCl, 1 mM EDTA, 0.5% IGEPAL CA-630 (Sigma-Aldrich #I8896-100ML), 0.25% Triton X-100, and 1X cOmplete protease inhibitor (Roche # 11873580001). After a 10-minute incubation with rotation at 4°C, nuclei were pelleted by centrifugation at 1000g and 4°C and the pellet was resuspended in 10 mL buffer containing 10 mM Tris-HCl pH 8.0, 200 mM NaCl, 1 mM EDTA, 0.5 mM EGTA, and 1X cOmplete protease inhibitor. After a 5-minute incubation with rotation at 4°C, nuclei were pelleted by centrifugation at 1000g and 4°C and the pellet was resuspended in 1 mL LB3 buffer containing 10 mM Tris-HCl pH 8.0, 100 mM NaCl, 1 mM EDTA, 0.5 mM EGTA, 1X cOmplete protease inhibitor, 0.5% N-laurylsarcosine, and 0.1% sodium deoxycholate and loaded in milliTUBE 1 ml AFA Fiber tubes (Covaris #520135). The chromatin was sonicated at the MIT BioMicro Center with a Covaris E220e sonicator using the following sonication conditions: 140W peak incident power, 5% duty factor, and 200 cycles per burst at 4°C for 20 minutes. Following sonication, 100 μl of LB3 with 10% Triton X-100 was added to each sample and the samples were centrifuged at 20,000g for 10 minutes at 4°C. The supernatant was reserved for the immunoprecipitation.

Prior to the immunoprecipitation, protein A Dynabeads (ThermoFisher #10001D) and protein G Dynabeads (ThermoFisher #10003D) were blocked. Beads were transferred to a magnetic rack, and the supernatant was removed. The beads were washed with 1 mL of ChIP dilution buffer containing 1% Triton X-100, 1 mM EDTA, 20 mM Tris-HCl pH 8.0, and 150 mM NaCl. The supernatant was removed, and the beads were resuspended in ChIP dilution buffer supplemented with 0.2 mg $mL^{-1}$ BSA (Sigma-Aldrich #B8667-1.25ML) and 0.05 mg $mL^{-1}$ yeast tRNA (ThermoFisher #AM7119) and incubated for 1 hour at 4°C with rotation. Following blocking, the beads were washed twice with 1 mL ChIP dilution buffer. 450 µg of chromatin was used as input to the immunoprecipitation. First, the chromatin was suspended in 1 mL ChIP dilution buffer and 30 µl of blocked Dynabeads was added. The chromatin was pre-cleared by rotation at 4°C for 2 hours. The Dynabeads were removed and 10% of the chromatin was reserved as input. The remaining chromatin was incubated overnight at 4°C with 15 µg of the following antibodies: CTCF (Millipore #07-729) or ZNF143 (Novus Biologicals H00007702-M01).

The next day, 10 µl of blocked Dynabeads was added to each sample and the samples were incubated at 4°C for 4 hours with rotation. Protein A beads were used for the CTCF immunoprecipitation and protein G beads were used for the ZFP143 immunoprecipitation. After removing the supernatant, the beads were washed by sequential 4 minute incubations at 4°C with 1 mL of the following buffers: low salt buffer (0.1% SDS, 0.1% Triton X-100, 2 mM EDTA, 20 mM Tris-HCl pH 8.0, and 150 mM NaCl), high salt buffer (0.1% SDS, 0.1% Triton X-100, 2 mM EDTA, 20 mM Tris-HCl pH 8.0, and 500 mM NaCl), LiCl buffer (250 mM LiCl, 0.1% IGEPAL CA-630, 10 mg $mL^{-1}$ sodium deoxycholate, 1 mM EDTA, and 10 mM Tris-HCl pH 8.0), and two washes with TE buffer. Following the washes, the beads were eluted in 100 µl of elution buffer containing 1% SDS and 100 mM $NaHCO_3$ by mixing at 25°C for 30 minutes at 1000 rpm. The supernatant was taken as the immunoprecipitated chromatin and incubated overnight at 65°C and 1000 rpm with 200 mM NaCl and 0.2 mg $mL^{-1}$ RNase A (ThermoFisher # EN0531). The following day, proteinase K (Viagen Biotech #501-PK) was added to a concentration of 0.2 mg $mL^{-1}$ and the samples were incubated for 1 hour at 45°C. The samples were purified using the Zymo ChIP DNA Clean and Concentrator kit (Zymo Research #D5205).

Prior to library preparation, the samples were transferred to Covaris microTUBE AFA Fiber Pre-Slit Snap-Cap tubes (Covaris #520045) and sonicated at the MIT BioMicro Center with a Covaris E220e sonicator using the following sonication conditions: 105W peak incident power, 5% duty factor, and 200 cycles per burst at 4°C for 50 seconds. The purified DNA proceeded to library preparation following the NEBNext Ultra II Library Prep kit (New England Biolabs #E7645L) with NEBNext Multiplex Oligos for Illumina (New England Biolabs # E7600S), and the libraries were purified by two sequential 0.9X Ampure XP bead purifications (Beckman Coulter #A63881). Library quality was assessed via Fragment Analyzer and qPCR at the MIT BioMicro Center. Biological duplicates were performed for all ChIP-seq libraries. Libraries were pooled in a 1:1 molar ratio based on the qPCR results and sequenced on a NovaSeq 6000 S2 flow cell at the Broad Institute of MIT and Harvard.

#### Precision Run-On Sequencing (PRO-seq) library preparation

Precision run-on sequencing (PRO-seq) libraries were prepared according to the method described by Mimoso and Goldman[89]. Briefly, 5 million cells were harvested for each condition and replicate and permeabilized in buffer containing 0.1% IGEPAL CA-630 (Sigma-Aldrich #I8896-100ML) and 0.05% Tween 20 (VWR #M147-1L). Cell permeabilization was confirmed by addition of trypan blue (ThermoFisher #15250061). The permeabilized samples were mixed with spike-in cells and equilibrated at room temperature for 5 minutes. The sample was then added to the run-on master mix containing the following components: 5 mM Tris-Cl, 5 mM $MgCl_2$, 0.5 mM DTT, 150 mM KCl, 10 µM of each biotin-11-NTP, 0.2 U $\mu l^{-1}$ SUPERase In (Invitrogen #AM2694), and 0.5% sarkosyl added fresh. The reaction was incubated at 37°C for 5 minutes and was stopped by addition of Buffer RL from the Norgen Total RNA Purification Kit (Norgen Biotek, #37500). RNA was purified using the Norgen Total RNA Purification Kit. The purified RNA was chemically fragmented by incubation for 5 minutes at 94°C in buffer containing 75 mM Tris-HCl pH 8.3, 112.5 mM KCl, and 4.5 mM $MgCl_2$. The samples were placed on ice and quenched by addition of 50 mM EDTA.

The samples were incubated with Dynabeads MyOne Streptavidin C1 (ThermoFisher #65001) at room temperature for 20 minutes. The beads were then washed twice with high-salt wash (50 mM Tris-HCl pH7.4, 2M NaCl, 0.5% Triton X-100, and 1 mM EDTA), twice with binding buffer (10 mM Tris-HCl pH 7.4, 300 mM NaCl,

0.1% Triton X-100, and 1 mM EDTA), and twice with low-salt wash (5 mM Tris-HCl pH 7.4, 0.1% Triton X-100, and 1 mM EDTA) all supplemented with SUPERase In. Beads were resuspended in TRIzol (Invitrogen #15596018), and the supernatant was pelleted by ethanol precipitation. The air-dried pellet was resuspended in water with 2 μM 3' adapter. The following ligation reaction was prepared and incubated overnight at 16°C: 15% PEG 8000, 1x T4 RNA ligase buffer, 0.5 U $\mu l^{-1}$ SUPERase In, and 10 units $\mu l^{-1}$ T4 RNA ligase 2, truncated KQ (New England Biolabs, #M0373). Samples were resuspended in binding buffer containing 1.25 M betaine (Sigma-Aldrich #61962) and heated for 5 minutes at 65°C. The bead binding was repeated as above and 5' decapping was performed using mRNA decapping enzyme (New England Biolabs, #M0608S). The samples were washed once each with high-salt wash, low-salt wash, and 1x PNK buffer (New England Biolabs, #M0201) and resuspended in PNK master mix containing T4 PNK (New England Biolabs, #M0201) and SUPERase In. Beads were washed with high-salt wash, low-salt wash, and 0.25x ligase buffer, and they were incubated for 2 hours at room temperature in mix containing 1x T4 RNA ligase 1 buffer, 1 mM ATP, 15% PEG 8000, 0.5 U $\mu l^{-1}$ SUPERase In, and 0.5 U $\mu l^{-1}$ T4 RNA ligase 1 (high concentration; New England Biolabs, #M0437). The RNA was converted to a cDNA library using SuperScript IV reverse transcriptase (Invitrogen, #18090200). Libraries were amplified using the NEBNext Ultra II Q5 Master Mix (New England Biolabs, #M0544) and cleaned up using the ProNex Size-Selective Purification System (Promega, #NG2001). Libraries were quantified by qPCR using the NEBNext Library Quant Kit for Illumina (New England Biolabs, #E7630), pooled in a 1:1 molar ratio based on the qPCR results, and sequenced on a NovaSeq 6000 at the Harvard FAS Bauer Core Facility. PRO-seq library construction, sequencing, and data analysis was performed by the Nascent Transcriptomics Core at Harvard Medical School, Boston, MA.

#### Fluorescence recovery after photobleaching (FRAP)

Prior to imaging, HaloTag cell lines were labeled with 500 μM Halo-JFX554 for 15 minutes followed by a 30-minute washout, and the $SNAP_f$-tag cell lines were labeled with 500 μM cpSNAP-JFX554 for 30 minutes followed by a 1-hour washout. The FRAP experiments were performed on a Zeiss LSM 900 Airyscan confocal microscope with a plan-apochromat, 63x, 1.40 NA, oil objective and an S2 DARK Premium humidified incubation chamber maintained at 37°C with 5.5% $CO_2$. The imaging was performed in confocal mode. For ZFP143 imaging, the following excitation parameters were used: 561 nm laser with 1.2% power, 53 μm pinhole size corresponding to 1AU, scan speed 6, and 800V detector gain. For imaging the H2B-Halo and Halo-NLS controls, the following parameters were used: 561nm laser with 0.2% power, 53 μm pinhole size corresponding to 1 AU, scan speed 6, and 700V detector gain. And for imaging the CTCF conditions, the following parameters were used: 561 nm laser with 1.0% power, 53μm pinhole size corresponding to 1 AU, scan speed 6, and 720V detector gain. The time series was acquired in a two-stage multi-acquisition. During the first stage, we acquired 30 frames with 1 second between frames, and during the second stage, we acquired 158 frames with 4 seconds between frames resulting in a total movie length of 662 seconds. A 1 μm circular spot was bleached in each movie after 10 frames using the following parameters: 561 nm laser with 100% power and 35 iterations per bleach at scan speed 7. Between 24 and 30 movies were acquired per condition across at least three different days.

#### Fast single-particle tracking (fast SPT)

Prior to imaging, cells were labeled at single-molecule density with either Halo-JFX554 for 15 minutes followed by a 30-minute washout. The following concentrations of dye were used: H2B-Halo and Halo-NLS 50 pM, clone D and C87 Halo-CTCF 100 pM, clones A and B ZFP143-Halo 500 pM, WT ZFP143-Halo and $DBD_{mut}$ ZFP143-Halo transfections 500pM, and $DBD_{null}$ ZFP143-Halo transfections 2.5nM. SPT experiments were performed on a custom-built Nikon Ti2-E microscope with an apochromat, 100x, 1.49 NA, oil objective, a perfect focus system, and a Prime 95B sCMOS camera (Teledyne Photometrics) with a 593/40 nm bandpass filter. The excitation laser angle was controlled with the iLas 2 motorized dual galvo system (Gataca Systems), and the sample conditions were humidity controlled and maintained at 37°C with 5% $CO_2$ with the Okolab stage top chamber. Samples were excited with a 1W 561 nm Coherent Genesis laser at 35% power modulated by an acousto-optic tunable filter (AA Opto-Electronic, AOTFnC-400.650-TN) triggered on the camera exposure TTL signal. The imaging ROI was set to a 200x200 pixel region around the nucleus, the camera exposure was set to 6 ms, and the excitation laser was pulsed for 1 ms during the camera exposure. One-minute-long movies were acquired, and all hardware was controlled through NIS-Elements software (Nikon).

#### Absolute abundance quantification

Absolute abundance quantification was performed as previously described[109,110]. Briefly, cells were labeled with 500 µM Halo-JFX554 for 30 minutes. Cells were subsequently washed three times with PBS and prepared for flow cytometry. They were then dissociated with TrypLE Express Enzyme, resuspended in PBS, and strained into Falcon™ Round-Bottom Polystyrene Test Tubes with Cell Strainer Snap Cap (Falcon, #352235) on ice. Cells were loaded onto a BD FACSCelesta Cell Analyzer (BD Biosciences) and 10,000 live cells were analyzed as determined by forward and side scatter gating. JFX554 fluorescence was excited using a 561 nm laser and read out using a 586/15 emission filter. A standard curve was then generated using previously determined CTCF abundances[109] for the C32 and C87 cell lines as well as wild type mESCs. The average abundance per cell of ZFP143 was taken as the average value of a line fit to the standard curve.

#### Analysis of Micro-C data

Paired-end sequencing reads generated by the Illumina NovaSeq sequencers were downloaded as .bcl files and converted to. fastq files using bcl2fastq (Illumina, v2.20.0.422). Read quality was verified using FastQC (v0.12.1), and reads were aligned to either the UCSC mm39 genome or the UCSC hg38 genome using bowtie2[111] (v2.2.5) with the following options: -local -reorder -very-sensitive-local. For consistency, the 150 bp reads from the NovaSeq X were trimmed to 66 bp, the NovaSeq S1 read length, using seqtk trimfq (v1.4). Aligned reads were then parsed using pairtools (v0.3.0) with the following command: pairtools parse --add-columns mapq --walks-policy mask --min-mapq 2 --drop-sam --drop-readid. Parsed reads were then sorted using pairtools sort, and pcr duplicates were removed using pairtools dedup with the --max-mismatch 1 and --mark-dups options. The reads were converted to a .cool format using cooler (v0.9.1) cload pairs and this was converted to a .mcool format with 10000000 5000000 2500000 1000000 500000 250000 100000 50000 25000 10000 5000 2000 1000 500 250 resolutions using cooler zoomify with the –balance option. For analyses involving the combined datasets across conditions, .cool files were merged using cooler merge before conversion to .mcool format.

Analysis was performed using custom python scripts (https://github.com/ahansenlab/ZNF143_analysis_code). For *de novo* loop calling using Mustache[112] (v1.3.1), the -pt 0.05 option was used, and loops were called at 10 kb, 5 kb, and 2 kb resolution. To create a consensus set of loops, loops called at all three resolutions were combined and for overlapping loops, the highest resolution call was taken. Loops overlapping features from ChIP-seq were identified using a previously reported custom script[87] (https://github.com/ahansenlab/RCMC_analysis_code/blob/main/loopFeatureOverlap.R) and can be found in **Tables S2-6.** Reproducibility matrices were generated using hicreppy[113] (v0.1.0). P(s) curves were computed using cooltools[114] (v0.5.4) expected_cis and derivative curves were computed using numpy (v1.23.5) gradient. Compartments were called at 500 kb resolution using cooltools eigs_cis and insulation scores were computed with 1 kb, 2 kb, 5 kb, 10 kb, 25 kb, and 50 kb windows using cooltools insulation. For insulation score metaplots, insulation scores were exported in .bigwig format and plotted using computeMatrix (v3.5.2) reference-point at boundaries as identified by cooltools. Aggregate peak analysis (APA)/loop pileup analysis was performed using coolpuppy[115] (v1.0.0) plotpup. Pileups were plotted as observed over expected as computed by cooltools expected_cis in log scale at 5000 bp resolution. Enrichment scores were calculated using the get_enrichment function from coolpuppy.lib.numutils. Region visualizations were generated using a combination of coolbox (v0.3.8) and pygenometracks (v3.8).

#### Analysis of ChIP-seq data

Paired-end sequencing reads generated by the Illumina NovaSeq sequencers were downloaded as .bcl files and converted to. fastq files using bcl2fastq (Illumina, v2.20.0.422). Read quality was verified using FastQC (v0.12.1), and reads were aligned to a concatenatated version of the UCSC mm39 genome and the UCSC hg38 genome using bowtie2[111] (v2.2.5) with the following options: --no-mixed --no-discordant. Aligned reads were sorted using sambamba (v0.6.6) and duplicates were removed with sambamba markdup. The reads aligning to the primary genome and the spike-in genome were separated using samtools (v1.6) view and converted to final

.bam format files using sambamba index. The bams were randomly downsampled with factors computed as the spike-in normalized ratio to the input ChIP-seq using sambamba. For visualization, the bams were converted to bedGraph format using macs2[116] (v2.2.6) pileup and subsequently to bigwig format using UCSC wigToBigWig (v2.8).

Analysis was performed using custom python scripts (https://github.com/ahansenlab/ZNF143_analysis_code). Reproducibility matrices were generated using multiBamSummary and visualized using plotCorrelation and plotPCA from deeptools[117] (v3.5.2). Peaks were called using macs2 callpeak with default values from the combined untreated files for both the ZFP143 ChIP-seq and the CTCF ChIP-seq. The called peaks can be found in **Tables S7-10**. ChIP-seq metaplots were generated using computeMatrix (v3.5.2) reference-point with the blacklist option using a blacklist from ENCODE[118]. Overlaps between peak sets, transcription start sites, or enhancer regions were calculated using bedtools[119] (v2.30.0) intersect. Similarly, bedtools intersect was used to identify enhancers as the regions with overlapping H3K4me1 and H3K27ac peaks in both mESCs and HEK293T cells. The reported enhancer regions can be found in **Tables S11-12**. To assign ZFP143 motifs to called peaks, instances of the ZNF143 motif from JASPAR were identified using fimo (v5.3.0) with the option --thresh 1e-3. Motif occurrences were converted to .bed format and the best scoring motif was kept for each called ZFP143 peak. To identify significantly changed ChIP-seq peaks between depletion conditions, we used a previously described[108] R script based on DEseq2[120] (v1.40.2). Spike-in read counts were used to calculate DESeq2 size factors used for normalization of the raw mm39 read counts, and significantly changed peaks (fold change > 1.5 and p-adj < 0.05) were plotted as an MA plot. Region visualizations were generated using a combination of coolbox (v0.3.8) and pygenometracks (v3.8).

#### Analysis of PRO-seq data

Basic analysis of PRO-seq data was conducted according to the method described by Mimoso and Goldman[89] using previously described custom scripts (https://github.com/AdelmanLab/NIH_scripts). Paired-end sequencing reads generated by the Illumina NovaSeq sequencers were downloaded as .bcl files and converted to. fastq files using bcl2fastq (Illumina, v2.20.0.422). Read pairs were trimmed using cutadapt (v1.14) with the options -O 1 –match-read-wildcards -m 26 and aligned to a concatenatated version of the UCSC mm39 genome and the UCSC dm6 genome using bowtie2. The spike-in and primary reads were separated and deduplicated using UMI-tools and separated using samtools (v1.3.1). These were subsequently converted to bedGraph format using a previously described custom script (bowtie2stdBedGraph.pl). For visualization, these were converted to bigwig format using UCSC wigToBigWig. Active transcription start sites were identified using a custom script (proTSSCall; https://github.com/NascentTranscriptionCore/proTSScall). Briefly, PRO-seq 3' read bedGraphs for plus and minus strands were separately combined across samples and the total read counts were assigned to the TSS plus 150 bp using the filtered TSS annotation described above. TSSs with less than or equal to 4 counts in this window are deemed 'inactive' and the remaining TSSs are collapsed to yield a single dominant TSS per gene. The dominant TSS is defined as the one with the highest TSS-proximal read count. If the highest read count is shared between multiple transcripts, the TSS furthest upstream, in a strand-aware fashion, is called dominant. Dominant TSSs sharing the same start position are deduplicated as follows: (1) if start positions are equal, the TSS with the longest associated annotated transcript is called dominant, (2) if start positions and transcript lengths are both equal, the TSS associated with the lowest Ensembl gene ID is dominant.

For differential expression analysis, reads were summed within the TSS to TES window for each active gene using the using the make_heatmap script (https://github.com/AdelmanLab/NIH_scripts), which counts each read one time, at the exact 3' end location of the nascent RNA. DEseq2, using the Wald test, was used to determine statistically significant differentially expressed genes, using the default size factors. The raw DESeq2 outputs can be found in **Tables S13-16**. The DESeq2 output was plotted as a volcano plot with a fold change threshold of 2 and a p-adjusted threshold of 0.001. Metaplots over gene bodies were generated using computeMatrix scale-regions. Gene ontology (GO) term enrichment analysis was conducted using a custom script based on the fgsea[121] (v1.26.0) R package using the mouse GO .gmt file (https://www.gsea-msigdb.org/gsea/msigdb/mouse/collections.jsp) (v2023.1).

#### FRAP analysis

FRAP analysis was performed in three steps. First, the movies were corrected for cell motion and drift, second the movies were photobleaching corrected and background subtracted, and third the aggregate tracks were fit and analyzed. To correct for movement of the bleach ROI, movies were loaded into a custom Python GUI (https://github.com/ahansenlab/ZNF143_analysis_code) and the position of the bleach ROI was manually updated to match the location of the bleach spot for each frame. The mean fluorescent intensity of the bleach, $I_{\text{bleach}}(t)$, was taken as the average intensity within the ROI. To estimate the average nuclear intensity, $I_{\text{nonbleach}}(t)$, and the average background intensity, $I_{\text{background}}(t)$, a second ROI and third ROI were drawn at representative locations within the nucleus and outside of the cell respectively. Cells with excessive drift or deformation of the bleach ROI were manually excluded from the analysis.

To correct for photobleaching, a series of correction factors ($C(t)$) were computed as the following ratio:

$$C(t) = \frac{I_{\text{nonbleach}}(t) - I_{\text{background}}(t)}{\langle I_{\text{nonbleach}}(t) - I_{\text{background}}(t)\rangle_{\text{prebleach}}}$$

Next, the FRAP curves were normalized according to the following equation:

$$\overline{I_{\text{bleach}}(t)} = \frac{C(t)(I_{\text{bleach}}(t) - I_{\text{background}}(t))}{\langle C(t)(I_{\text{bleach}}(t) - I_{\text{background}}(t))\rangle_{\text{prebleach}}}$$

where $\overline{I_{\text{bleach}}(t)}$ is the normalized, background-subtracted, and photobleaching corrected FRAP signal.

Finally, the following reaction-dominant model[84] was fit to the average of $\overline{I_{\text{bleach}}(t)}$ across all movies in each condition:

$$FRAP(t) = 1 - C_{\text{eq,fast}}e^{-k_{\text{off,fast}}t} - C_{\text{eq,slow}}e^{-k_{\text{off,slow}}t}$$

where $k_{\text{off,slow}} < k_{\text{off,fast}}$.

#### SPT Analysis

All SPT movies were tracked using the version of the Hungarian algorithm implemented in quot[92] (https://github.com/alecheckert/quot/tree/master). The exact parameters used for tracking are given here: https://github.com/ahansenlab/ZNF143_analysis_code/blob/main/imaging_analysis/tracking_config.toml. The tracks were then limited to only those within the nucleus using a custom nuclear masking script (https://github.com/ahansenlab/ZNF143_analysis_code). For Spot-On[86] fitting of the jump length distributions, the python implementation of Spot-On (https://gitlab.com/tjian-darzacq-lab/Spot-On) was used with the following parameters: cdf=True, use_entire_traj=True, frac_bound=[0, 1], d_free=[0.15, 25], d_bound=[0.0005, 0.08], sigma_bound = [0.005, 0.1], iterations=3, dZ=0.700. The localization error was determined to be 22.3 nm by fitting to the H2B-Halo condition and was input as a parameter to the remaining sets. To generate state arrays of diffusion coefficients, saSPT[92] (https://saspt.readthedocs.io/en/latest/, v0.4.0) was used with the following parameters: likelihood_type = RBME, focal_depth = 0.7.
